# Assessing Microcirculation Impairment in Ischemic Stroke Mice Using Arteriovenous Co-fluctuation Analysis

**DOI:** 10.64898/2026.07.08.737374

**Authors:** Bochao Niu, Yanlin Bi, Benjamin Klugah-Brown, Yang Yuan, Quandan Tan, Guoliang Zhu, Yapeng Lin, Junli Hao, Kejie Chen, Lingling Wang, Zhe Kang Law, Hongyan Gong, Jie Yang

## Abstract

Accurate assessment of cerebral hemodynamics impairment traditionally relies on arterial metrics, yet often overlooks venous drainage and arteriovenous dynamics, thereby limiting the evaluation of ischemia-induced microvascular dysfunction. To address this limitation, we implemented a signal-averaging framework, combined with co-fluctuation analysis, to extract predominantly arterial and venous hemodynamic signals and construct a dynamic arteriovenous co-fluctuation index that quantifies frame-by-frame coordination between arterial inflow and venous outflow activity. This time-resolved index enables spatial characterization of large-scale cortical arteriovenous coordination beyond conventional static correlation-based analyses. Comparative analyses between healthy controls and acute ischemic stroke mice demonstrated that the arteriovenous co-fluctuation index sensitively detects disruption of vascular coordination, revealing a slower state transition that occurs alongside distinct temporal abnormalities and regional heterogeneity between ischemic core and penumbral regions. These findings underscore the utility of arteriovenous coordination as a sensitive indicator of microcirculatory dysfunction, offering a practical analytical tool for assessing stroke-induced microvascular impairment.

## Introduction

Recanalization therapies such as intravenous thrombolysis and endovascular mechanical thrombectomy are effective treatments for patients with large vessel occlusions (Jia et al., 2024; Liu et al., 2026; Sperring et al., 2023). However, a substantial proportion of patients continue to exhibit persistent neurological impairment despite successful macrovascular recanalization, a phenomenon commonly referred to as “no-reflow” (Jia et al., 2024; Sperring et al., 2023). This clinical dissociation indicates that restoration of large-vessel patency alone is insufficient to fully recover downstream microvascular perfusion and tissue viability. Preservation of ischemic penumbral tissue depends on coordinated function across the arterial-capillary-venous axis, requiring not only adequate arterial inflow but also effective venous drainage and the maintenance of capillary-level perfusion dynamics.

Current clinical perfusion imaging approaches, including CT perfusion, MR perfusion, Doppler ultrasound, and positron emission tomography, primarily emphasize arterial perfusion-related parameters such as cerebral blood flow (CBF) (Kropf et al., 2025; Leigh et al., 2018), cerebral blood volume (CBV) (Lakhani et al., 2025; Picchi et al., 2025), and mean transit time (MTT) (Bulut et al., 2022; Hilkens et al., 2024; Leigh et al., 2018). Although these metrics have substantially advanced stroke diagnosis and therapeutic decision-making, they remain predominantly focused on arterial supply and provide limited characterization of venous drainage dynamics and integrated arteriovenous coordination. Consequently, conventional perfusion metrics may incompletely capture tissue-level vascular dysfunction, particularly in regions where impaired venous outflow or disrupted vascular synchronization contributes to ongoing hypoperfusion despite preserved arterial inflow (Li et al., 2023; Proner et al., 2026).

Ischemic stroke initiates a multiscale cascade of hemodynamic disturbances that progressively disrupt vascular coordination across both macrovascular and microvascular systems. At the macrovascular level, collateral circulation is rapidly recruited through redistribution of blood flow via pial anastomoses and leptomeningeal vessels (Glück et al., 2024). Our prior work demonstrated enhanced functional integration efficiency within whole-brain networks of the ischemic hemisphere (Niu, Wu, et al., 2024), suggesting that large-scale vascular compensation may be accompanied by adaptive network-level reorganization. At the microvascular level, tissue perfusion becomes impaired through mechanisms including capillary constriction, pericyte dysfunction, and increased vascular resistance (Sun et al., 2021; Xu et al., 2025). Importantly, our prior studies further demonstrated disruption of the normal temporal synchrony between arterial inflow and venous outflow dynamics following ischemia (Niu, Sihai, et al., 2024), representing a fundamental breakdown in arteriovenous coordination that extends beyond simple flow reduction. These observations collectively demonstrate that the functional outcome of microcirculation in ischemia is determined by integrated dysfunction across the arterial-capillary-venous axis, wherein impairment at any level compromises the viability of the entire microvascular unit.

Despite increasing recognition of vascular coordination in cerebrovascular physiology, current in vivo optical imaging approaches remain limited in their ability to characterize dynamic interactions between arterial and venous hemodynamics. Although high-resolution imaging techniques can quantify parameters such as capillary red blood cell velocity, perfused capillary density, and tissue blood volume (Gong et al., 2025; B. X. Liu et al., 2025; Wang et al., 2021; Xu et al., 2025), these approaches typically generate composite hemodynamic signals arising from sequential vascular compartments, including arteries, capillaries, and veins. This spatial overlap makes selective isolation of arterial and venous hemodynamic signatures particularly challenging in large-scale cortical imaging datasets.

Several technical factors contribute to this limitation. First, the dense spatial interdigitation of cortical arteries and veins creates substantial anatomical overlap, complicating compartment-specific signal segregation (Park et al., 2023; Schmid et al., 2019). Second, because optical imaging methods detect hemodynamic activity through erythrocyte-derived contrast signals, both arterial and venous compartments contribute strong signal intensity, particularly in regions where vessels are closely apposed (Boas & Dunn, 2010; Han et al., 2026). In addition, local blood flow directionality and velocity profiles may not differ sufficiently to permit reliable computational separation in all cortical regions. Third, the limited depth resolution of optical imaging systems introduces out-of-focus scattering and signal contamination from vessels located at different cortical depths (Schmid et al., 2019). As a result, superficial veins and deeper penetrating arterioles may contribute overlapping signals that are difficult to reliably separate using conventional post-hoc.

Consequently, venous drainage dynamics and coordinated arteriovenous interactions remain underrepresented in conventional cerebrovascular analyses. To address this limitation, we developed a co-fluctuation-based analytical framework to characterize the dynamic coordination between arterial and venous blood flow activity. Inspired by the theoretical framework of Esfahlani et al. (Zamani Esfahlani et al., 2020), demonstrating that static functional connectivity can be decomposed into frame-wise co-fluctuation events, we reformulated negative arteriovenous correlations as temporally evolving anti-phase fluctuation dynamics. By computing instantaneous products of normalized arterial and venous signals across time, we generated continuous arteriovenous co-fluctuation time series that transform conventional static correlation measures into a dynamic index of vascular coordination. This framework enables time-resolved assessment of arterial-venous synchronization and provides a systems-level approach for investigating vascular dysfunction in ischemic stroke beyond traditional artery-centric perfusion models.

## Materials and Method

### 2.1 Experimental Animal Preparations

All animal procedures adhered to ARRIVE guidelines and complied with the ethical standards of the Laboratory Animal Welfare Ethics Committee of Qingdao University, under approval number 202301C575420240908. To verify the method’s robustness and applicability, we conducted evaluations on multiple datasets comprising both rat and mouse models. One group of adult male Sprague-Dawley rats (3–4 months, 280–320 g, n = 19) was sourced from our prior research (Niu, 2026). Another group of rats (6–8 months, 300–350 g, n = 30) was obtained from previous work (Niu et al., 2026). Adult male C57BL/6J mice (8–12 weeks, 20–25 g, n = 30 for control group, n = 19 for stroke group) were obtained from Qingdao Daren Fucheng Animal Co., Ltd. Animals were individually housed in specific pathogen-free conditions, with ad libitum access to food and water, under a 12-hour light/dark cycle. For surgical preparation, animals were anesthetized using isoflurane (4% for induction, 1.5%–2.0% for maintenance) and positioned in a stereotaxic frame. Core body temperature was monitored via rectal probe and maintained at 37±0.5°C using a feedback-controlled heating system. Ophthalmic ointment was applied to prevent corneal drying during the procedure.

Whole-brain laser speckle contrast imaging was performed using a thinned-skull preparation. Briefly, the scalp was incised along the midline, and the periosteum was bluntly dissected to expose the skull. The bone surface was cleaned with 3% hydrogen peroxide, then gradually thinned using a spherical drill under continuous saline irrigation to achieve optical transparency while minimizing thermal damage.

Postoperatively, animals received subcutaneous carprofen (5 mg/kg) for postoperative analgesia and were housed in a temperature-controlled recovery chamber at 30°C for at least 24 hours before imaging. All surgical and imaging procedures were performed by experienced personnel to ensure animal welfare and experimental consistency.

### 2.2 Resting-State Imaging for Cortical Cerebral Blood Flow

Cortical cerebral blood flow was monitored using laser speckle contrast imaging (LSCI) systems (RFLSI III, RWD Life Science, China, for mice; SIM BFI HR Pro, SIM Opto-Technology, China, for rats). Both systems utilized the safety threshold (< 20 mW/cm²) to ensure adequate tissue penetration without thermal damage. Imaging parameters were optimized for whole-cortex coverage, with pixel resolutions of 549 × 600 (∼14.6 μm/pixel) for mice and 494 × 656 (42.5 × 44.2 μm²) for rats, at sampling rates of 5 Hz and 10 Hz, respectively.

Resting-state CBF data were acquired under stable isoflurane anesthesia from the following rodents: two groups of Sprague-Dawley rats (n = 19 and n = 30), and one group of C57BL/6J mice (n = 30). Animals were securely immobilized in a stereotaxic frame to minimize motion artifacts, and 10-minute baseline recordings were obtained for each animal. All imaging sessions were conducted in a controlled environment that was maintained at a constant temperature, sound-isolated, and light-shielded. Core body temperature was kept at 37±0.5°C via a feedback-regulated heating platform throughout data acquisition to preserve the physiological stability of cerebral perfusion.

### 2.3 Induced MCAO Model Preparation

Focal cerebral ischemia was modeled using transient middle cerebral artery occlusion (tMCAO) in male C57BL/6J mice (8–12 weeks of age, n = 19). Animals were anesthetized with 2% isoflurane, and a silicone-coated nylon filament (0.23 mm tip diameter) was advanced 9–11 mm along the external carotid artery to occlude the left middle cerebral artery origin, as in our previous study (Gong et al., 2025). After 60 minutes of occlusion, the filament was withdrawn to reperfuse blood flow. Core body temperature was maintained at 37°C during surgery and for 2 hours postoperatively to minimize temperature-related variations in ischemic injury.

Resting-state CBF imaging was performed in each animal within 72 hours following tMCAO induction. All imaging procedures, acquisition parameters, and physiological monitoring protocols were identical to those used for healthy control mice.

### 2.4 Signal Preprocessing Pipeline

Resting-state CBF imaging was preprocessed using a standardized pipeline developed and validated in our research (Niu et al., 2026; Niu, Wu, et al., 2024). The preprocessing workflow began with spatial normalization: cortical spatial masks were generated based on stereotaxic coordinates from rat and mouse brain atlases, and affine registration was applied to align individual CBF maps to a standard rat brain template. Individualized masks were then applied to each CBF image to exclude non-cerebral background and environmental noise.

The CBF time series was preprocessed via four sequential steps: **(1) Denoising:** Raw RGB images were converted to gray images and smoothed twice with Gaussian filters. The first pass used a 5 × 5 pixel kernel with a standard deviation of 1.3 to suppress mechanical noise (White et al., 2011), the second employed a Gaussian kernel with a full width at half maximum of 5 pixels to enhance the signal-to-noise ratio through spatial smoothing (Bergonzi et al., 2015). (2) **Head motion regression:** Head motion parameter was extracted from the background activity and regressed out from the blood flow signal to minimize motion-related artifacts. **(3) Outlier correction:** Temporal outliers in the time series were corrected using nearest-neighbor interpolation. **(4) Detrend:** Linear trends and very slow oscillations (< 0.01 Hz) were removed from the time series to eliminate slow signal drift.

### 2.5 Vascular Mask Construction Framework

A critical limitation of optical imaging is its limited depth of field (L. J. Liu et al., 2025; Park et al., 2023), which prevents clear discrimination between arteries and veins at different cortical depths (**Fig. 1A**). To address this constraint, we developed a comprehensive segmentation framework to construct reliable arterial and venous masks, integrating multiple strategies across spatial scales and anatomical regions to ensure methodological robustness, anatomical accuracy, and broad applicability.

**Fig. 1.**
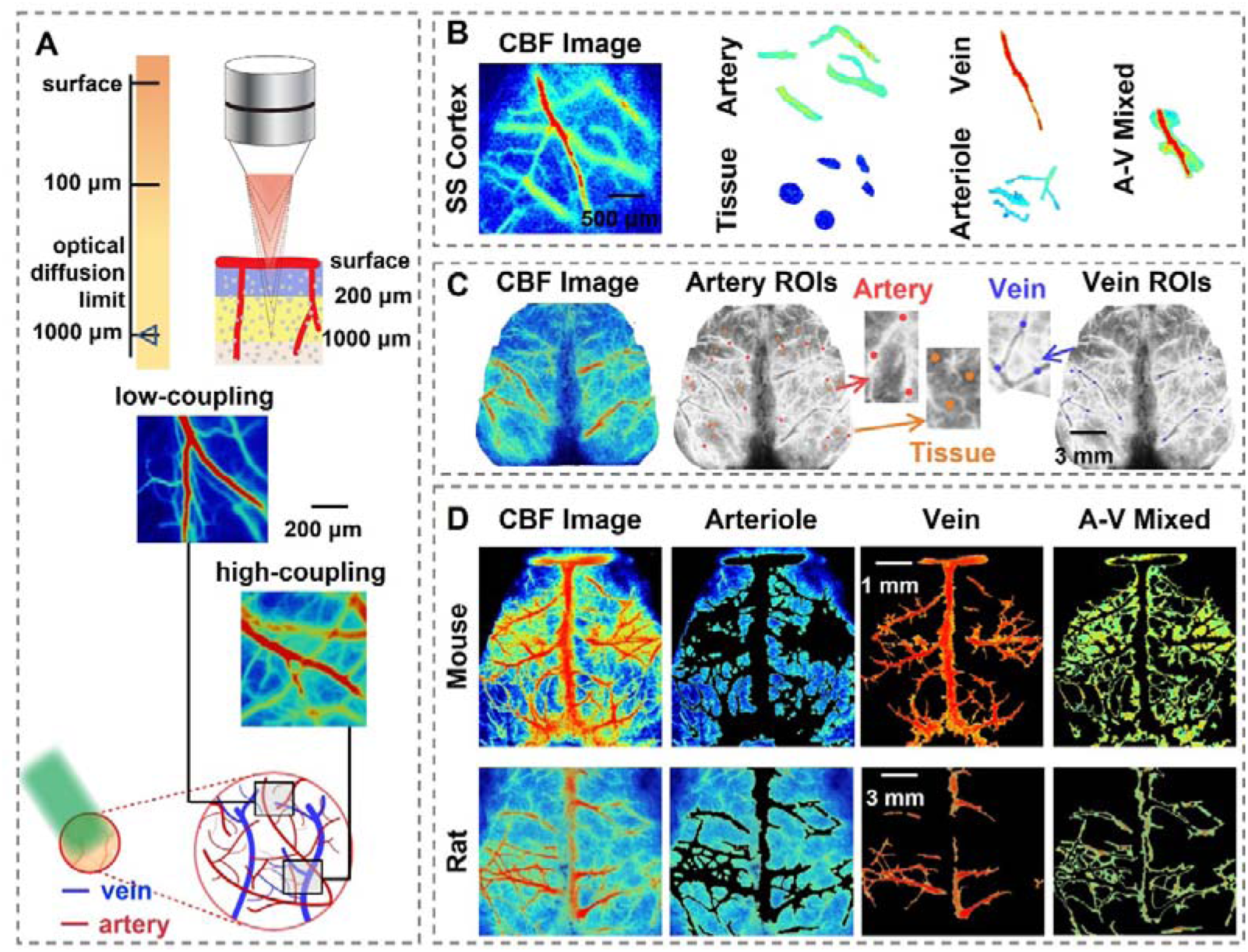
Segmentation masks of different vessel types from raw CBF images. **A.** Arterial and venous vessels are tightly coupled in the cerebral cortex. **B.** Manually segmented masks for large arteries, arterioles, cerebral tissue, veins, and mixed arteriovenous regions. **C.** Arterial, tissue, and venous masks from manual ROI placement. **D.** Masks of arteries, veins, and mixed arteriovenous regions derived from k-means clustering segmentation.

#### Strategy 1: high-resolution manual segmentation

In the rats’ somatosensory-motor cortex (n = 19), which is densely vascularized, five distinct compartments were manually segmented based on morphological criteria: large arteries, small arteries, brain parenchyma, veins, and mixed arteriovenous regions (**Figs. 1B, S1**). This labor-intensive approach generates anatomically faithful reference masks with precise boundary definition, serving as a benchmark for automated segmentation validation. However, the extensive manual effort limits its wide application across entire brain volumes.

#### Strategy 2: representative ROI placement

Multiple ROIs in the healthy mice (n = 30) were selected from arterial trunks, venous trunks, and non-vascular brain parenchyma (**Figs. 1C, 4A**), serving as representative samples of arterial, venous, and background signals, respectively. This sampling strategy provides operational simplicity and broad applicability with minimal computational burden. Its primary limitation lies in the potential failure to capture whole-brain hemodynamic variability when spatial sampling density is insufficient.

#### Strategy 3: semi-automated clustering-based segmentation

K-means clustering (k = 5) was applied to the denoised CBF maps of both rats and mice (n = 30 for each species) to partition the image space into distinct signal domains. Temporal correlation matrices were computed across cluster-derived signals, with negatively correlated clusters identified as arterial and venous masks (**Fig. 1D**). The remaining clusters were categorized as mixed vascular regions or background noise based on spatial distribution. This semi-automated workflow enables efficient whole-brain segmentation; cluster boundaries may require post-hoc refinement guided by the arteriovenous anticorrelation principle.

### 2.6 Validation of Signal Purification Robustness

To verify the reliability of our spatial averaging strategy for arterial and venous signal purification, we conducted a comprehensive robustness assessment through three complementary analyses:

#### (1) Signal homogeneity within vascular masks

Using manually segmented arterial, venous, and mixed vascular masks, we computed functional connectivity matrices from all pixels within each mask. Distribution characteristics of correlation coefficients, including kurtosis and skewness, were analyzed to quantify signal heterogeneity. Distributions exhibiting high kurtosis and positive skewness reflected synchronized blood flow activity and minimal compositional heterogeneity, whereas deviations from this pattern suggested potential high heterogeneity. In addition, the proportion of negative correlations within each matrix was quantified to measure cross-contamination between arterial and venous compartments.

#### (2) Stability of the spatial averaging procedure

Bootstrap resampling was performed on individual vascular masks at progressively increasing sampling densities (ranging from 5% to 95% of the total number of pixels) to generate random subsets of the original signal population. For each subset, the averaged time series was compared against the full-mask reference using Pearson correlation and phase difference. High similarity at low sampling rates demonstrated the stability of the spatial averaging approach.

#### (3) Quantifying resilience to segmentation errors

To assess robustness against realistic segmentation errors, we synthetically mixed arterial mask signals with increasing proportions of venous signals (0.05 to 0.50 in 0.05 steps), generating a continuum of hybrid signal profiles. The correlation between each hybrid signal and the original arterial signal was tracked across mixing levels. The mixing threshold was defined as the proportion at which correlation dropped to 90% of its initial value, thereby quantifying the maximum tolerable venous interference for reliable signal purification.

### 2.7 Methodological Framework for Dynamic Arteriovenous Coupling Analysis

Recent advances by Esfahlani and colleagues (Zamani Esfahlani et al., 2020) established a mathematical insight showing that static functional connectivity emerges from the temporal integration of instantaneous co-fluctuation patterns across brain networks. This insight has prompted a paradigm shift in neuroscience, moving the field from static to dynamic analyses of brain connectivity. The co-fluctuation approach quantifies the instantaneous synchronization of hemodynamic signals between brain regions via edge-time series analysis. Its temporal average is mathematically equivalent to the Pearson correlation, thereby providing a rigorous mathematical foundation for reinterpreting static connectivity within a dynamic framework.

Our prior investigation revealed that arterial and venous blood flow signals in the cerebral cortex exhibit anti-phase oscillations and negative correlations (Niu, Sihai, et al., 2024). This observation challenges traditional interpretations of arteriovenous relationships, suggesting that negative correlations reflect not merely static statistical properties, but rather the cumulative outcome of discrete, transient anti-phase fluctuation events occurring throughout the hemodynamic time series. Guided by this co-fluctuation perspective, we developed a novel framework for constructing arteriovenous co-fluctuation time series, transforming the Pearson negative correlation into a frame-by-frame arteriovenous circulation index. This approach enables direct quantification of coupling dynamics between arterial inflow and venous outflow, allowing real-time tracking of microvascular coordination. The methodology comprises three integrated components, as shown in Figure 2:

**Fig. 2.**
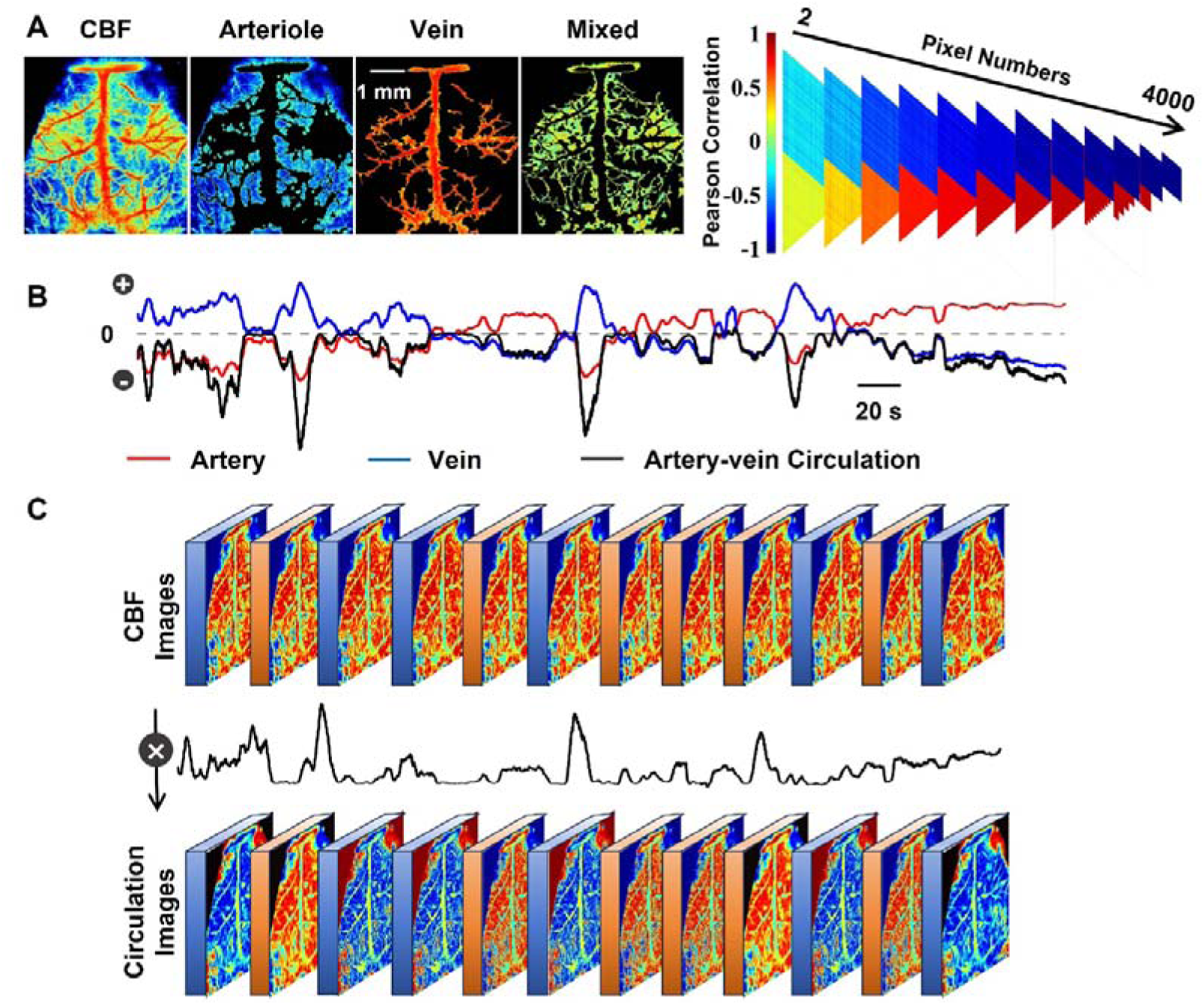
Computational pipeline of arteriovenous fluctuation coupling. **A.** Vascular masks extracted from clustering maps were used for signal purification via pixel averaging. Purified arterial and venous signals showed a strong negative correlation. **B, C.** Purified arterial and venous signals were multiplied to yield a negative arteriovenous circulation time series. This time series was then converted to positive values and utilized to amplify the raw cerebral blood flow time series, thereby enabling real-time whole-brain circulation mapping.

**Step 1: Vascular blood signal purification.** We generated mean time series for the arterial and venous compartments by spatially averaging blood flow signals across all voxels within their respective vascular masks. This spatial averaging effectively suppresses local noise while preserving the dominant oscillatory characteristics of each vascular compartment.

**Step 2: Instantaneous arteriovenous circulation index computation.** The extracted time series were Z-score normalized to eliminate scale differences and facilitate direct comparison between arterial and venous signals. Recognizing the inherent anti-oscillation relationship between arterial inflow and venous outflow during the resting state, we developed a specialized transformation to address the challenge posed by negative product values, which can potentially obscure strong coupling states.

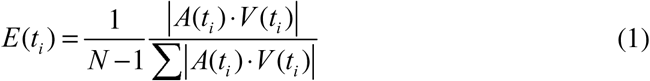

where *i* indicates each time point, *A*(*t*) and *V* (*t*) represent the normalized arterial and venous time series. Specifically, we converted the instantaneous product sequence into a probability density function by: (1) computing the absolute area under the product curve, and (2) normalizing by the total integrated area. This transformation yields a non-negative coupling metric, *E*(*t*), where higher values indicate stronger moment-to-moment arteriovenous synchronization.

**Step 3: Spatiotemporal retransforming of whole-brain microvascular dynamics.** We applied the time-varying coupling index as a dynamic weighting factor to the original resting-state CBF time series through frame-wise multiplication. This generates a spatiotemporal map of arteriovenous coupling-modulated hemodynamic activity, enabling comprehensive characterization of microvascular coordination dynamics across the entire brain.

### 2.8 Characterizing Spatiotemporal Dynamics of Microcirculatory Activity

The co-fluctuation analysis framework was used to dynamically characterize cerebral blood flow microcirculatory states through integrated temporal and spatial pattern analysis, aiming to elucidate microcirculatory functional impairments induced by stroke.

Given that temporal co-fluctuation amplitudes exhibited a right-skewed distribution, prompting us to define the top 5% of timepoints based on amplitude distribution as high-coupling states, and the bottom 5% as low-coupling states. The former represents optimal microcirculatory coordination, and the latter reflecting poorest coordination. Principal component analysis was applied to timepoints corresponding to these two states, with spatial activity patterns derived from the first principal component to compare regional blood flow distribution between high and low coupling conditions. To further evaluate stroke-induced alterations in microcirculatory dynamic transitions, arteriovenous time series from both stroke and control groups were normalized and combined for k-means clustering (cluster number ranging from 3 to 6) to identify shared functional states. State-specific metrics, including temporal occupancy, duration, occurrence times, self-transition probability, and between-state transition probabilities, were employed to compare groups and quantitatively characterize stroke-related abnormalities in microcirculatory state transitions.

To reveal spatial patterns of CBF and microcirculatory impairment, we used the contralateral hemisphere as an internal reference to quantify ischemic injury severity. For each region in the ischemic hemisphere, we calculated the difference in signal intensity relative to its contralateral homolog as a measure of regional damage. Based on established criteria from previous studies (Biose et al., 2020; C. Li et al., 2022; Li et al., 2013), regions below 70% of the contralateral value were classified as the ischemic core, and those with 50–70% of the contralateral intensity were classified as the ischemic penumbra. Intra-regional homogeneity was assessed using Pearson’s correlation to quantify regional coherence, complemented by phase difference analysis to evaluate regional synchronization. Finally, we examined associations between cortical vascular composition and ischemic vulnerability indicators including ischemic volume and penumbra/core ratio. Correlation analyses were performed, with significance assessed via FDR-corrected P-values (q < 0.01) and model fitting evaluated using correlation coefficients.

### 2.9 Statistical Analysis

Statistical analyses were conducted in MATLAB. Data normality was assessed using the Kolmogorov-Smirnov test and Q-Q plots. For comparisons among multiple groups, one-way ANOVA was performed. For parameter comparisons of different vascular types obtained by manual segmentation, the Mann-Whitney U test was applied. For pairwise comparisons, two-tailed paired t-tests (n = 30) compared differences between top and bottom states within the same animal group. Two-tailed unpaired t-tests were used to compare microcirculation parameters between healthy controls and mice with induced ischemic stroke. To control for multiple comparisons, false discovery rate corrections were implemented. Statistical significance was set at *p < 0.05, **p < 0.01, and ***p < 0.001.

### 2.10 Large Language Models Statement

The authors declare that Qianwen AI large model was employed only for linguistic refinement and readability enhancement of the manuscript. No AI tools were involved in study design, data collection/analysis, result interpretation, or the drafting of scientific arguments and conclusions.

## Results

### 3.1 Characterization of Cerebral Blood Flow in Masks of Different Vasculatures

In vivo optical imaging of cerebral hemodynamics faces a fundamental resolution constraint: the limited optical depth-of-field precludes clear spatial separation of closely positioned arteries and veins at different cortical depths (Glück et al., 2024; Park et al., 2023). When optical blur and light scatter cause signal overlap between vessels at distinct depths, mathematical isolation of pure arterial or venous components becomes technically infeasible, even with post-hoc signal processing (**Fig. 1A**). Current research usually overlooks this limitation (Fischer et al., 2023; Mo et al., 2023; Wang et al., 2024; Wang et al., 2021), either treating mixed vascular signals as homogeneous or focusing exclusively on arterial dynamics, thereby hindering a penetrating understanding of arteriovenous coordination mechanisms. We therefore hypothesize that optical imaging captures composite signals from anatomically stacked arterial and venous structures. These vascular networks exhibit region-specific structural coupling patterns: highly coupled regions prevent signal separation while loosely coupled regions allow partial discrimination. Despite being invisible in raw imaging datasets, this anatomical coupling mechanism exerts a profound influence on the accurate interpretation of CBF dynamics.

To validate this hypothesis, we employed a manual segmentation strategy to construct five distinct vascular masks (**Fig. 1B**) that characterize region-specific vascular architecture. In high-coupling regions where arterial and venous networks are densely interwoven, we delineated three separate vascular compartments: venous-dominant, arterial-dominant, and mixed arteriovenous territories. In contrast, in low-coupling regions characterized by spatially segregated vascular networks, we focused on two functionally distinct tissue types: small arteriolar beds and adjacent parenchymal tissue. The segmented vascular masks and statistical information are presented in Supplementary Figure S1 and Supplementary Table 1, respectively. These precisely segmented masks enabled the extraction of resting-state CBF signals, facilitating the analysis of spatial variations in arteriovenous coupling patterns across cortical regions. We then validated these functional data using two perspectives: global signal properties and local spatial homogeneity, revealing how anatomical coupling strength modulates the dynamic patterns of CBF activity.

First, we averaged all pixel signals within each vascular mask to generate a mean time series and compared the differences among these time series. Spectral power analysis revealed that in the 0.01–0.2 Hz band, the power proportion in veins (0.58±0.16) was significantly lower than that in brain tissue (0.66±0.15, P_FDR_ = 0.006). In the 0.2–0.74 Hz band, arteries exhibited the highest power proportion (**Fig. 3A**), whereas arterioles and brain tissue showed the lowest. In addition, signals from the same vessel type exhibited strong positive correlations (**Fig. 3B**) with near-zero time lags. In contrast, heterogeneous signals between arteries and veins showed strong negative correlations with larger time lags.

**Fig. 3.**
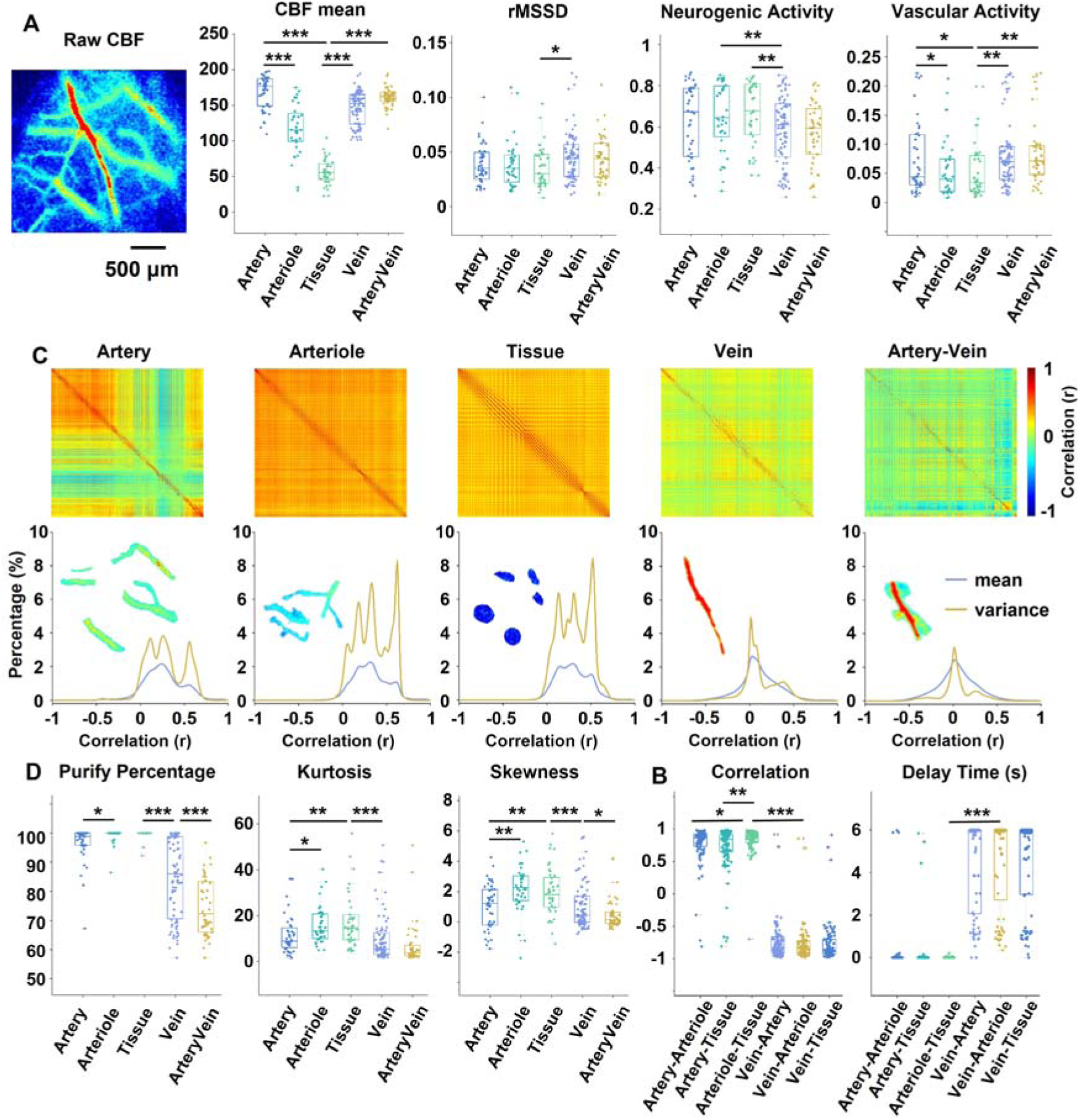
Signal quality comparison across different vascular masks. **A.** Arteries exhibit the highest cerebral blood flow and the highest percentage of vascular activity power (0.2–0.74 Hz), whereas cerebral tissue shows the opposite trend. **B.** Signals from homogeneous arteries show a strong positive correlation with a time delay near 0 s. In contrast, signals between arteries and veins are negatively correlated with a larger time delay. **C, D.** Arterioles and cerebral tissue display the strongest internal similarity in correlation matrices, along with the highest kurtosis and skewness, suggesting more consistent signal characteristics in these regions. Metric differences across vascular components are assessed using the Mann–Whitney U test with FDR adjustment. Detailed statistics for each vasculature type are provided in Table S1. *p < 0.05, **p < 0.01, ***p < 0.001

We then evaluated pixel-level homogeneity within each vascular mask using Pearson’s correlation analysis. Among the five vascular masks, the correlation matrices of arterioles and brain tissue showed the highest homogeneity (**Figs. 3C, S2**), followed by arteries, and the arteriovenous mixed regions exhibited the lowest homogeneity. Quantitative analysis further confirmed that the arteriole and brain tissue masks had the highest pixel purity (**Fig. 3D**), whereas arteriovenous mixed regions had the lowest. Analysis of kurtosis and skewness demonstrated that correlation matrices in arterioles and brain tissue exhibited higher kurtosis and skewness (e.g., arterioles kurtosis 15.9±8.4 vs. arteries kurtosis 11.4±8.1, P_FDR_ < 0.001).

Collectively, these findings reveal significant spatial heterogeneity in arteriovenous vascular coupling, which is closely associated with resting-state blood flow dynamics. In high-coupling regions characterized by dense arterial-venous interweaving, signal contamination prevents the independent discrimination between arterial and venous signals. By contrast, in low-coupling regions where vascular networks are relatively segregated, vascular masks of small arteriolar beds and parenchymal tissue display high signal separation fidelity and enhanced internal homogeneity.

### 3.2 Blood Flow Signal Purification and Noise Effects on the Purification Process

In the previous section, we demonstrated significant spatial heterogeneity in arteriovenous coupling across cortical regions. In high-coupling regions, tightly interwoven arterial and venous networks hinder signal separation. In low-coupling regions, vascular signals separate more clearly with better internal consistency within masks. However, these observations merely characterize structural-functional relationships and do not yield practical methods for the complete separation and purification of arterial and venous signals. This methodological gap limits both the accurate assessment of venous drainage and a deeper understanding of arteriovenous coordination mechanisms. To address this limitation, we implemented a manual ROI-based approach to define five distinct vascular compartments (**Fig. 4A**): arteries, veins, brain tissue, and their intersection regions. By quantifying how non-target vascular signals interfere with signal purification within each mask, we established a generalizable framework for arteriovenous signal separation and purification.

**Fig. 4.**
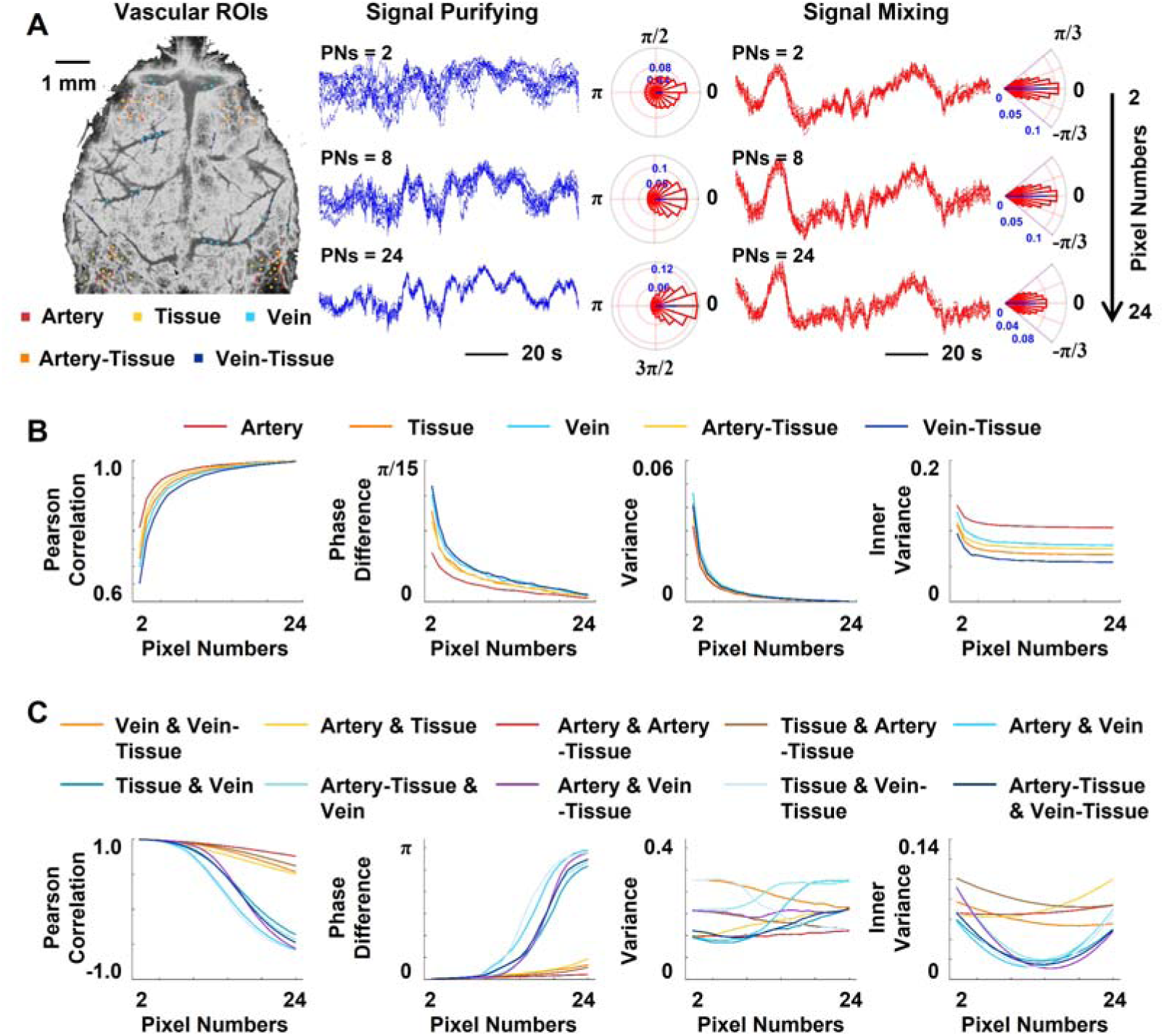
The illustration of signal purification and mixing across various vascular ROIs. **A.** As pixel count in ROIs increases, random CBF fluctuations diminish and phase differences tend toward 0°. However, when arterial ROIs incorporate venous signals, the phase deviation from 0° grows as the noise component increases. **B.** Using larger pixel counts for purification enhances the Pearson correlation with the reference, decreases phase differences, and reduces variance. **C.** Homogeneous mixing, such as tissue or artery–tissue, maintains the purified signal’s features, whereas heterogeneous venous mixing substantially changes these characteristics. For instance, a 50% arterial–venous mixture results in minimal correlation with the reference.

To isolate pure blood flow signals, we implemented a signal averaging technique and evaluated its performance through two complementary approaches. First, we examined signal convergence characteristics by progressively expanding ROI sizes and quantifying the technique’s efficacy in extracting pure vascular signals from mixed inputs. Second, we assessed noise resilience by measuring interference caused by both homogeneous vascular noise (originating from the same vessel type) and heterogeneous vascular noise (from different vessel types), thereby establishing the method’s performance boundaries under distinct noise conditions.

First, we manually selected ROIs to quantify blood flow signals from five distinct vascular compartments in the cerebral cortex (**Fig. 4A**): arteries, brain tissue, veins, artery-tissue junctions, and vein-tissue junctions. For each ROI, we performed signal averaging on randomly selected pixel subsets of varying sizes, using the mean signal from all pixels serving as the reference standard. Our results demonstrated that signal purification from small pixel subsets exhibits large variability around the reference signal (**Figs. 4A, S3A**), characterized by dispersed phase differences and low Pearson correlation coefficients. In contrast, when averaging more than half of the pixels within an ROI, signal fluctuations diminish markedly, phase differences converge toward zero, and correlation coefficients exceed 0.8. Further quantitative analysis reveals a nonlinear convergence pattern (**Figs. 4B, S4**), where averaging only 50% of the pixels achieved 92% similarity to the full ROI signal. This finding indicates substantial signal redundancy within vascular ROIs, suggesting that inherent signal consistency provides robust resilience against random fluctuations and heterogeneous noise during purification processes.

To systematically evaluate the impact of noise on signal purification, we conducted controlled simulations with noise components introduced quantitatively. Our analysis revealed distinct responses to different noise types. Homogeneous noise mixtures, such as arterial signals combined with parenchymal tissue or venous signals mixed with venous-parenchymal interface signals, exhibited minimal interference with purification performance (**Figs. 4C, S5–S6**). Remarkably, even when homogeneous noise constituted more than 50% of the total signal, purified outputs maintained > 95% similarity to reference standards. In contrast, heterogeneous noise mixtures, particularly arterial-venous signal combinations, induced a rapid performance degradation (**Figs. 4C, S5**), with similarity approaching zero when noise pixels surpassed the 50% threshold.

To precisely quantify heterogeneous noise effects, we implemented simulations using vascular masks generated through k-means segmentation. These results demonstrated that mask size fundamentally determines noise tolerance limits (**Figs. 5, S7–S9**). Using arterial masks as a representative case: with only 10 pixels, 9% venous noise contamination reduced similarity to below 90% of baseline values; however, expanding the mask to 50 pixels increased the noise tolerance threshold to 24%. Importantly, we identified a threshold effect governing purification efficacy: beyond 38% noise contamination, increasing the number of pixels provided no additional purification benefit, with similarity remaining below 90% of its initial value regardless of mask size.

**Fig. 5.**
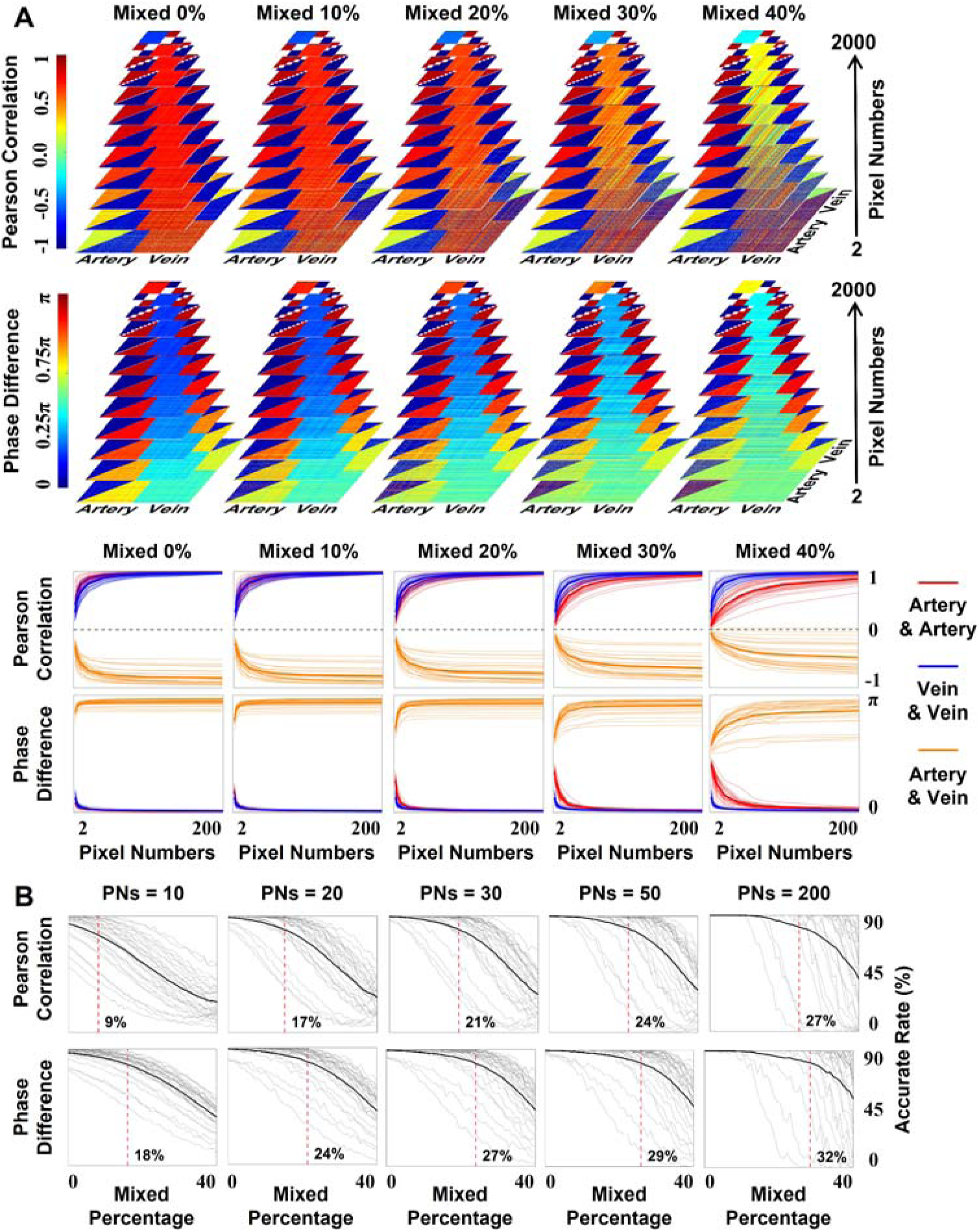
Impact of venous noise on arterial signal purification. **A.** When the venous noise ratio remains below the 20% threshold, arterial signal purity is maintained, as evidenced by intra-arterial correlation values approaching 1 and phase differences near 0°. In contrast, a 40% venous noise ratio exceeds the purification capacity, resulting in ineffective signal purification. **B.** As venous noise increases, the proportion of pixels exhibiting strong negative artery-vein correlations (r < −0.4) declines. With a low pixel count for purification, a 10% noise level reduces this proportion to 90% of its initial value. However, higher pixel counts enhance noise tolerance, delaying this decline.

### 3.3 Spatiotemporal Decoding of Whole-Brain Arteriovenous Microcirculation

Cerebral blood flow delivers essential oxygen and metabolic substrates to support neuronal function, making accurate measurement of its dynamics is critical for understanding brain physiology. However, optical imaging faces a persistent challenge: arterial and venous signals on the cortical surface cannot be fully separated (Glück et al., 2024; Park et al., 2023), severely compromising measurements of microcirculation. In our previous work, we mitigated this limitation by extracting venous signals and applying regression analysis to suppress their contamination (Niu et al., 2026; Niu, Wu, et al., 2024). Although this approach enhanced signal purity, residual venous influence persisted. Our early analysis revealed a key physiological insight: during resting states, arterial and venous blood flow signals exhibit robust anti-phase oscillations with strong negative temporal correlation (Niu, Sihai, et al., 2024). This intrinsic vascular coordination offers a natural basis for signal separation. Leveraging this insight, we developed a co-fluctuation analysis framework that transforms the arterial-venous negative correlation coefficient into a frame-by-frame vascular circulation index. Multiplying this index by the original CBF signal effectively isolates microcirculatory activity, enabling precise characterization of cortical hemodynamics.

Temporal analysis of whole-brain arteriovenous coupling signals revealed a strongly right-skewed distribution (**Fig. 6A**), characterized by intermittent bursting activity with prolonged periods at minimal baseline levels punctuated by brief, high-amplitude peaks. To investigate these contrasting microcirculatory states, we extracted the top 5% of timepoints (signal peaks) and the bottom 5% (troughs) for comparative analysis (**Fig. 6B, C**). Whole-brain activity patterns during peak moments closely resembled the mean signal across the entire recording period (r = 0.98±0.017), while exhibiting a strong negative correlation with trough timepoints (r = −0.89±0.35). Principal component analysis of whole-brain functional activity demonstrated remarkable spatial stability across temporal states (**Fig. 6C**), revealing consistent regional specificity with maximal activity in sensorimotor cortices and minimal engagement of temporal and limbic regions. Notably, peak moments showed the lowest variance explained by the first principal component (80.8%±12.2%), consistent with the rapid dynamic transitions observed at these timepoints. To evaluate regional differences in hemodynamic variability, we compared microcirculatory dynamics across arterial, venous, and arteriovenous mixed vascular territories. Arteriovenous mixed regions exhibited significantly higher temporal variability than other vascular compartments (**Fig. 6D**). Cross-species validation revealed that rats exhibited microcirculatory activity patterns highly consistent with those observed in mice (**Fig. S10**).

**Fig. 6.**
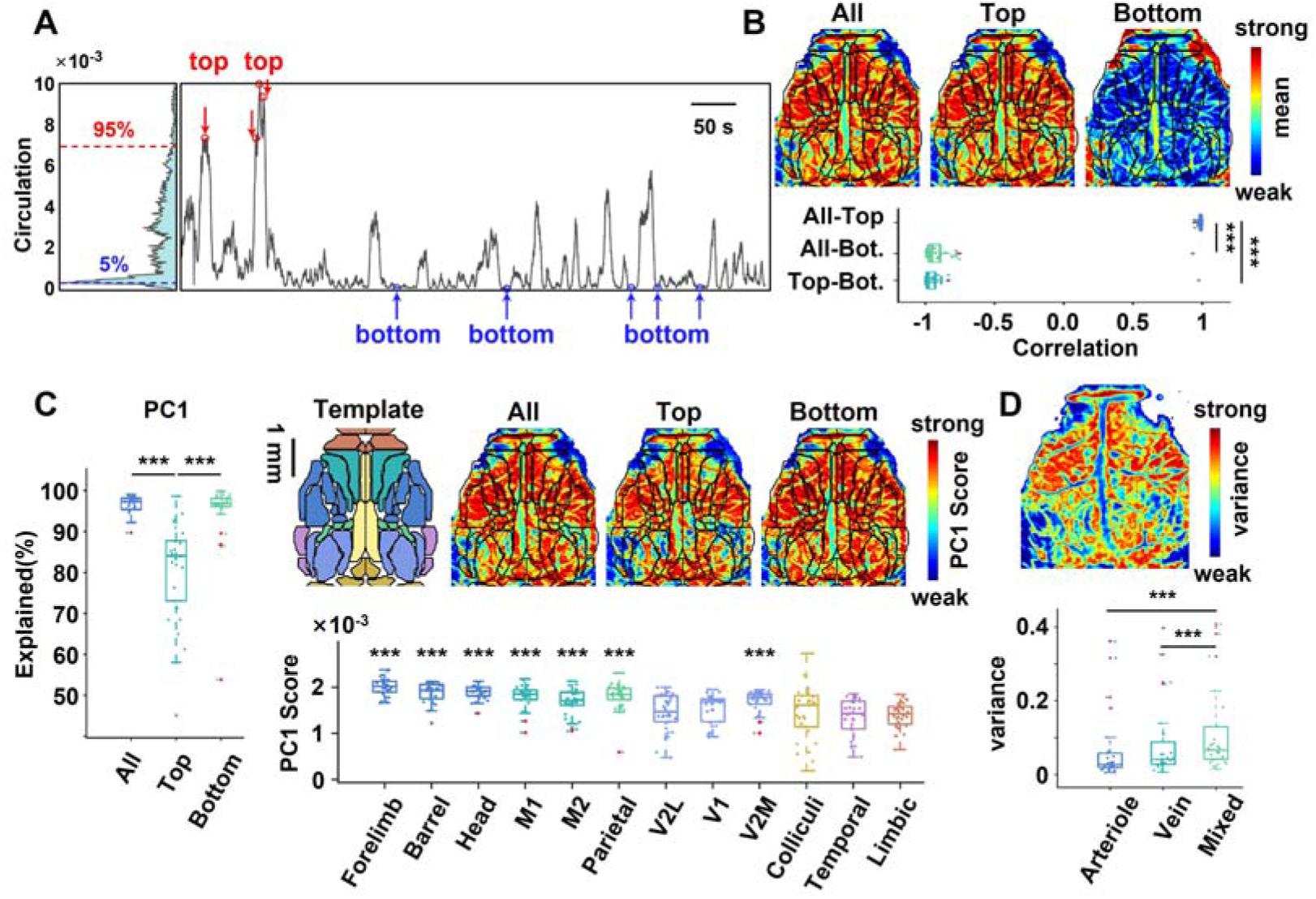
Spatiotemporal characteristics of whole-cerebral blood circulation activity in rats. **A, B.** The time series of arteriovenous fluctuation coupling exhibits a right-skewed distribution. Timepoints corresponding to the top 5% (peaks) and bottom 5% (troughs) of this distribution were extracted for further analysis. The cortical activity pattern at peaks closely resembles that of the full time series, but is negatively correlated with that at troughs. **C.** The spatial pattern of cortical circulation activity is highly consistent across peak, trough, and full signals, with maximal activity in the sensorimotor cortex and minimal activity in the temporal and limbic cortices. Interregional parameter differences are examined by one-way ANOVA (n = 30), followed by pairwise t-tests with FDR correction using V2L as the reference. **D.** Arterial and venous regions showed lower signal variance than arteriovenous mixed regions. Comparisons of individual items across timepoints and vascular components are assessed via paired t-test with FDR adjustment (n = 30). ***p < 0.001 in B, C, and D, *** indicates p < 0.001 whose PC1 scores are greater than secondary visual cortex in C. Bot.: bottom

To assess whether cerebral blood flow exhibits a consistent spatial organization analogous to microcirculatory dynamics, we analyzed CBF dynamics during the peak, trough, and entire signal periods within the circulation time series. The result showed that minimal spatial correlation between these different temporal states (r < 0.30). Principal component analysis of the CBF signal failed to identify a dominant spatial pattern (**Figs. S11, 12**), with no significant regional topological distinctions observed across the brain (P_FDR_ > 0.05).

### 3.4 Stroke-induced Impairment of Cerebral Microcirculatory Activity

In previous sections, we established that arteriovenous coordination serves as an effective biomarker of microcirculatory integrity, with significant spatial differences in microcirculatory activity observed across brain regions. These findings imply that a comprehensive assessment of post-stroke tissue viability requires evaluating both arterial inflow and venous outflow dynamics. We therefore hypothesized that analyzing arteriovenous coordination would provide superior detection of microcirculatory impairment compared to conventional arterial perfusion metrics alone. To test this hypothesis, we developed an acute ischemic stroke model, acquired resting-state functional CBF data, and conducted integrated analyses of temporal dynamics and spatial distribution patterns to assess the diagnostic efficacy of this approach.

Temporal dynamics analysis revealed significant alterations in arteriovenous coordination after ischemic stroke. K-means clustering of arteriovenous temporal profiles showed markedly reduced fluctuation velocity in stroke animals compared to controls (**Fig. 7A**), with probability density distributions showing an increased proportion of signal concentration within baseline minimum ranges (**Fig. 7B**). These findings corresponded to impaired state transition capacity in the stroke group. Although the relative proportions of functional states remained stable across different cluster numbers (k = 3 to 5) and their spatial activity patterns exhibited high similarity (**Figs. 7A, S13**), stroke animals displayed abnormal state persistence, characterized by increased intra-state retention probabilities (**Figs. 7C, S14**) and prolonged dwell times (**Fig. 7D**). Specifically, for k = 3 clustering, the self-transition probability for state 1 was significantly higher in stroke animals (0.98±0.009) than in controls (0.97±0.014; P_FDR_ = 0.0097). Conversely, inter-state transition probabilities were substantially reduced in stroke animals (**Figs. 7C, S15**), and transition times between distinct states were decreased (**Fig. 7D**). For example, the transition probability from state 1 to state 3 was significantly lower in stroke animals (0.014±0.009) compared to controls (0.024±0.013; P_FDR_ = 0.0096).

**Fig. 7.**
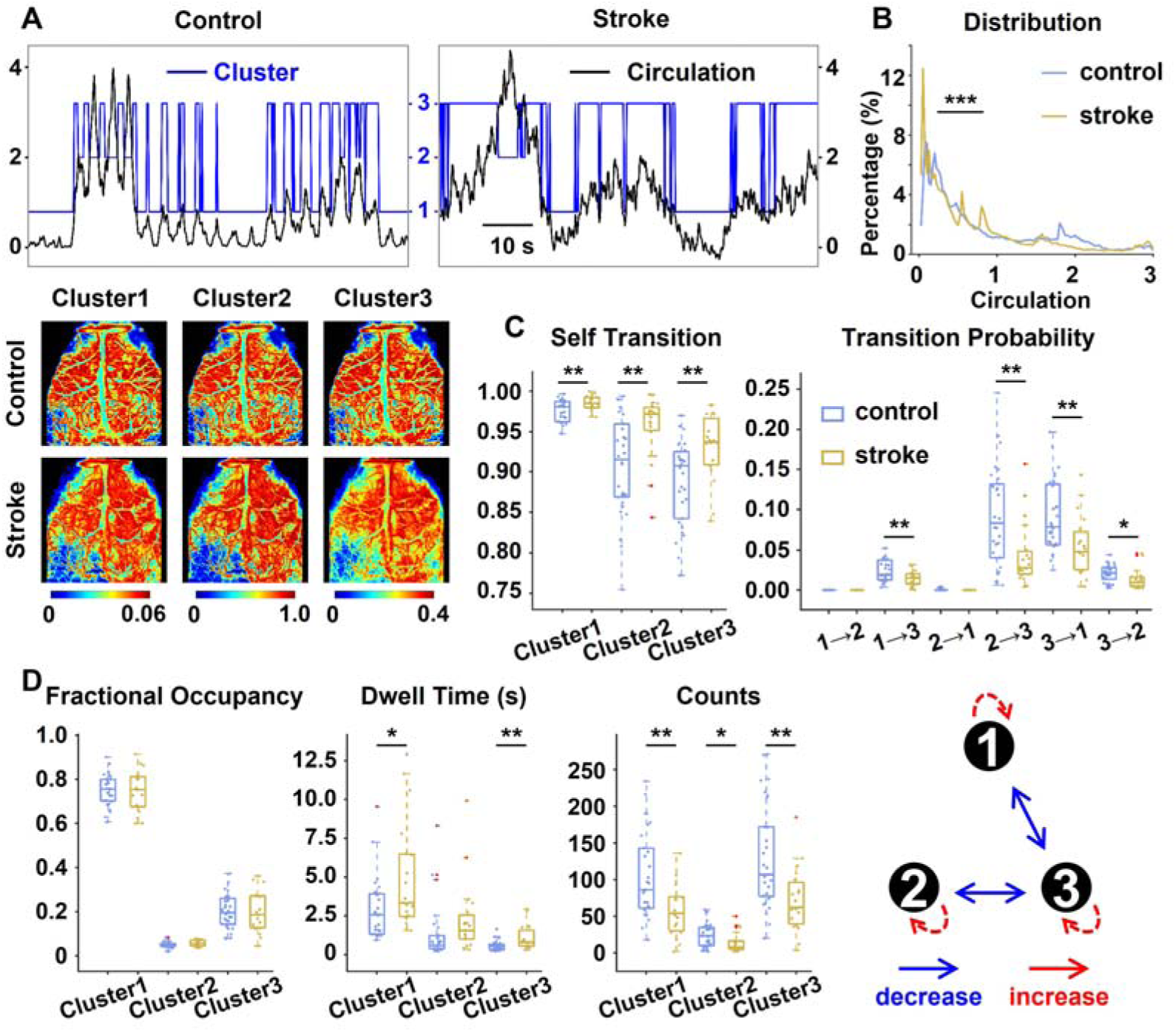
Stroke disrupts dynamic functional transitions in cerebral microcirculation. **A, B.** Stroke attenuates fluctuation dynamics in arteriovenous circulation. Compared to controls, stroke mice exhibit reduced signal fluctuation speed, increased right-skewness in probability density distributions, and increased signal occupancy within the minimum baseline range. **C, D.** Stroke impairs the capacity for functional state transition across three distinct microcirculatory states. Post-stroke microcirculation exhibits prolonged dwell times per state, with increased state-maintenance probability, while the inter-state transition probability is markedly reduced, ultimately decreasing overall state-transition events. Group differences between control (n = 30) and stroke (n = 19) are assessed using an unpaired t-test. *p < 0.05, **p < 0.01, ***p < 0.001

To clarify the spatial heterogeneity of microcirculation within the ischemic hemisphere, we performed a whole-brain analysis of microcirculation activity in the stroke model. Regions with lower mixed arteriovenous vasculature, specifically visual, auditory, and limbic motor cortices, exhibited severe microcirculatory impairment (**Fig. 8A**) and reduced CBF perfusion (**Fig. S16A**). In contrast, somatosensory-motor regions, characterized by a higher proportion of mixed arteriovenous vasculature, showed indistinguishable impairment (P_FDR_ > 0.05). This differential vulnerability prompted us to examine microcirculatory heterogeneity within the ischemic territory itself. Using established thresholds, microcirculatory signal intensity <30% of contralateral homologous regions for infarct core, and 30–50% for ischemic penumbra, we identified distinct functional signatures between these two territories. The penumbra exhibited greater temporal signal variability (**Fig. 8B**) but higher spatial homogeneity (**Fig. 8C**) than the ischemic core. Specifically, compared with ischemic core, functional connectivity matrices derived from penumbral pixels showed higher mean correlation coefficients (0.94±0.047 vs. 0.89±0.075; P = 0.003), smaller mean phase differences (0.051±0.047 vs. 0.081±0.082; P = 0.022), and lower inter-element variance within the matrices. Parallel analyses of CBF dynamics corroborated these findings (**Fig. S16**). Moreover, the ischemic vulnerable regions delineated by the microcirculation framework showed excellent spatial concordance with those identified by the CBF method (**Fig. S17**). Crucially, the microcirculation-based segmentation achieved a significantly greater statistical difference (T = 4.27) in the Pearson correlation between the ischemic penumbra and infarct core than the standard CBF approach (T = 2.53).

**Fig. 8.**
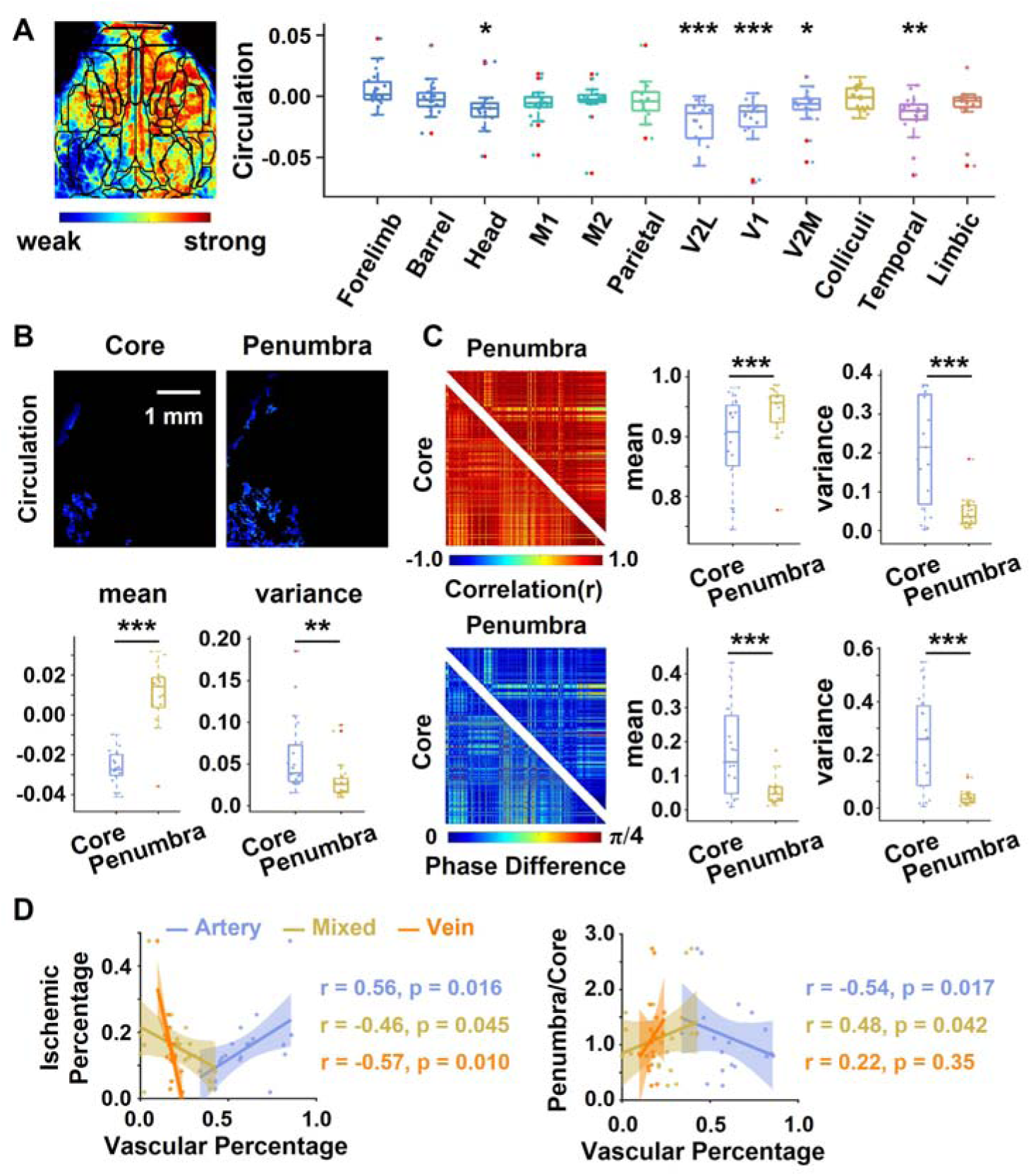
Quantitative profiling of microcirculatory impairment in ischemic regions. **A.** Microcirculation intensity is lower in the left visual and temporal regions compared to their right-hemisphere counterparts. Homologous differences between ischemic and contralateral hemispheres are analyzed with a oneLJsample tLJtest (n = 19). **B, C.** The ischemic penumbra exhibits preserved microcirculatory function relative to the impaired infarct core. Microcirculation signal intensity is significantly higher in the penumbra, accompanied by robust functional connectivity metrics, including higher mean inter-voxel correlation coefficients, smaller mean phase differences, and lower variance in functional connectivity matrices. Metric differences between the ischemic core and penumbra are assessed using a paired t-test (n = 19). **D.** Vascular architecture influences regional susceptibility to ischemic damage. Arterial vessel proportion is positively correlated with ischemic lesion size, while the proportions of arteriovenous mixed and venous vessels show inverse correlations. Of note, a larger fraction of mixed vascular regions predicts a higher penumbra-to-core ratio. *p < 0.05, **p < 0.01, ***p < 0.001

The regional heterogeneity in arteriovenous coupling across the brain prompted us to investigate whether vascular composition influences ischemic vulnerability. Our analysis revealed that arterial proportion was positively correlated with total ischemic lesion extent (rLJ=LJ0.56, P_FDR_ LJ=LJ0.016), whereas venous proportion (rLJ=LJ−0.57, P_FDR_LJ=LJ0.010) and arteriovenous mixed-region proportion (rLJ=LJ−0.46, P_FDR_ LJ=LJ0.045) both exhibited significant inverse correlations with lesion extent. However, lesion size alone provides an incomplete assessment of ischemic severity. Given the divergent tissue fates of ischemic penumbra and infarct core, we further examined how vascular composition relates to the penumbra/core ratio. Notably, arterial vessel proportion correlated negatively with this ratio (rLJ=LJ−0.54, P_FDR_ LJ=LJ0.017), while arteriovenous mixed-region proportion showed a positive correlation (rLJ=LJ0.48, P_FDR_ LJ=LJ0.042). Consistent with these observations, parallel analyses based on conventional CBF signals corroborated the close link between vascular composition and ischemic vulnerability (**Fig. S16D**).

## Discussion

In this study, we applied a co-fluctuation analysis framework to transform the inverse temporal relationship between arterial and venous hemodynamic activity into a frame-by-frame arteriovenous circulation index for assessing cerebral microcirculatory dynamics. Methodological validation demonstrated that both arterial and venous vascular masks contain heterogeneous signal components; however, spatial signal averaging effectively suppressed this contamination and enabled robust extraction of predominantly arterial and venous hemodynamic time series. This approach permitted characterization of large-scale arteriovenous coordination and construction of an arteriovenous circulation coefficient reflecting dynamic vascular synchronization. Under physiological conditions, cortical microcirculatory activity exhibited a conserved spatial organization, with strong activity observed in sensorimotor cortices and comparatively lower activity in temporal and limbic regions. In the ischemic stroke model, regions exhibiting stronger arteriovenous coordination demonstrated relative resistance to ischemic injury, whereas regions with weaker coupling suffered more severe impairment. By comparison, the ischemic penumbra maintained a more preserved microcirculatory and perfusion function than the infarct core, which was severely impaired. Temporal analyses further revealed that ischemic stroke disrupted normal microcirculatory dynamics, characterized by prolonged persistence within individual functional states and reduced inter-state transition probability.

### 4.1 Physiological Basis of Arteriovenous Oscillatory Coupling

The cerebral circulation functions as an integrated physiological network that coordinates arterial inflow, capillary-level exchange, and venous drainage to maintain metabolic homeostasis. Conventional neuroimaging approaches have largely examined arterial and venous compartments independently (Fu et al., 2024; L. J. Liu et al., 2025; Oelschlegel et al., 2024; Taxt et al., 2021), thereby overlooking their dynamic physiological interdependence. Although these approaches have substantially advanced our understanding of cerebral perfusion, they remain limited in their ability to characterize real-time, system-level vascular coordination. To address this limitation, we developed a co-fluctuation-based analytical framework that transforms the inverse temporal relationship between arterial and venous signals into a dynamic arteriovenous circulation index capable of tracking large-scale vascular synchronization over time.

Consistent with our previous studies (Niu et al., 2026; Niu, Sihai, et al., 2024), arterial and venous signals exhibited robust anti-phase oscillatory behavior under physiological conditions, reflected by negative Pearson correlations. Following acute ischemia, this relationship became substantially attenuated and, in some regions, shifted toward positive correlation (Niu, Sihai, et al., 2024), indicating disruption of normal vascular coordination. This anti-phase blood flow pattern is validated by multiple experimental findings (Glück et al., 2024; Liu et al., 2021; Shen et al., 2012; Skøtt et al., 2025). Fluorescent labeling studies (Liu et al., 2021) confirmed the opposing temporal fluctuation between arterial inflow and venous outflow signals. Furthermore, phase analysis (Glück et al., 2024) revealed that healthy cerebral vasculature maintains an arterial-venous phase difference exceeding 120°, which diminishes to approximately 60° during acute ischemic episodes. Collectively, these observations establish that the anti-phase relationship between arterial and venous flow is a fundamental characteristic of normal cerebral perfusion and a sensitive biomarker of impaired vascular coordination after ischemic injury. Importantly, while anti-phase synchronization likely reflects coordinated inflow-outflow dynamics, its precise physiological determinants remain incompletely understood and may involve contributions from vascular compliance (Lecordier et al., 2023; Xu et al., 2025), neurovascular coupling (Evans et al., 2025), transit delays (Glück et al., 2024), vasomotor oscillations (Li et al., 2024), and systemic physiological rhythms (Zhang et al., 2025).

### 4.2 Methodological Validation for Separating Arterial and Venous Signals

To ensure accurate separation of arterial and venous blood flow signals, we conducted a comprehensive validation of our signal separation methodology. In optical imaging, the spatial interweaving of vascular networks frequently results in individual pixels containing mixed arterial and venous components. Direct analysis of raw signals, therefore, introduces substantial confounding effects arising from overlapping flow dynamics, potentially compromising subsequent arteriovenous coupling analyses. We therefore rigorously evaluated our separation framework to determine whether the extracted arterial and venous signals reliably reflected their dominant hemodynamic profiles, thereby establishing a methodological basis for subsequent analyses of arteriovenous coordination.

To assess the feasibility of arterial-venous signal separation, we manually segmented cortical vascular masks. Because the sensorimotor cortex exhibits particularly high vascular density, five vascular compartments were defined: artery-dominant, vein-dominant, arteriolar, parenchymal tissue, and arteriovenous mixed regions. Quantitative analysis based on signal decomposition demonstrated that compartment-specific signal contribution was below 85% in both venous and arteriovenous mixed regions, whereas arteriolar and parenchymal tissue regions exhibited substantially higher signal specificity. These findings indicate intrinsic limitations in completely separating arterial and venous signals within highly interwoven vascular territories. The close spatial proximity of cortical vascular structures, together with their tight metabolic and hemodynamic coupling, constrains signal isolation (Grandjean et al., 2023; Xiong et al., 2017). This observation is consistent with previous neurovascular imaging studies demonstrating that highly interconnected vascular regions present substantial challenges for compartment-specific signal extraction (Park et al., 2023; Schmid et al., 2019).

Based on these findings, we developed an optimized framework combining ROI placement and semi-automated vascular segmentation to improve arterial-venous signal fidelity across the cortical surface. Notably, signal analysis demonstrated that using only 50% of the arterial ROI pixels preserved approximately 92% of the signal similarity relative to the full ROI, indicating functional redundancy in vascular masks that suppresses heterogeneous signal contamination. To quantitatively assess the robustness of this signal purification strategy, we conducted simulation experiments using mixed artery-vein signal processing. Simulation analyses further demonstrated that when spatial sampling exceeded 200 pixels, the correlation between purified signals and the original simulated source signals remained above 0.90 despite up to 38% heterogeneous signal contamination. These findings indicate that spatial averaging substantially suppresses stochastic noise and enhances the consistency of compartment-specific signal components.

Collectively, these results demonstrate that signal averaging effectively reduces heterogeneous noise and improves intra-compartment signal coherence, thereby enabling robust differentiation between arterial and venous hemodynamic activity. Our previous investigations similarly demonstrated that whole-brain signal averaging using limited pixel counts yielded relatively weak correlations, whereas increasing pixel numbers strengthened signal consistency (Niu, Sihai, et al., 2024). This principle is well supported in the literature. Bauernfeind et al. (Brigadoi et al., 2014) demonstrated that spatial averaging of multi-channel fNIRS signals significantly attenuates physiological noise, with noise reduction proportional to the square root of the number of averaged channels, thereby improving signal-to-noise ratio. Similarly, Zheng et al. (Zheng et al., 2019) showed that resting-state spatial filtering approaches, including common average referencing, enhance the stability and consistency of target neural activity signals. Together, these findings support the technical feasibility and physiological relevance of our signal purification strategy for dynamic hemodynamic analysis

### 4.3 Paradigm Shift in the Measurement of Blood Flow Microvasculature

The validated signal purification approach provides a methodological foundation for reframing the study of cerebral microcirculation. Rather than conceptualizing cerebral blood flow as a static transport system, our findings support the interpretation that cerebral vascular organization behaves as a dynamically coordinated network. Within this framework, arteriovenous synchronization may represent an emergent systems-level property reflecting coordinated vascular regulation across the arterial-capillary-venous axis.

For decades, research on blood flow dynamics has centered on static vascular metrics (Boas & Dunn, 2010; Jin et al., 2024; Xu et al., 2025), such as cerebral blood volume, flow velocity, and vessel caliber, by extracting time-averaged parameters from observation windows. These approaches have provided foundational insights for both basic neuroscience and clinical applications forward; they remain widely employed in studies of static functional connectivity (Aykan et al., 2025; Niu, Sihai, et al., 2024; Zhang & Jiang, 2025), amplitude of low-frequency fluctuations (Lu et al., 2024), and power spectral analysis (Niu et al., 2026). Yet this averaging approach ignores the rich temporal activity inherent to resting-state brain activity. Emerging evidence (Gutierrez-Barragan et al., 2022; Klugah-Brown et al., 2026; Lee et al., 2024; Yang et al., 2023) challenges the assumption of fixed temporal organization within brain networks. Instead, real-time coordination within brain networks exhibits aperiodic, spontaneous rhythmic characteristics (L. Li et al., 2022; Zamani Esfahlani et al., 2020). These time-varying characteristics unlock unique neural fingerprints that are specific to individual subjects (Lee et al., 2024; Yang et al., 2023; Zamani Esfahlani et al., 2020), offering a promising pathway toward personalized neurofunctional profiling.

In addition, our results provide compelling evidence for the fundamental principle that cerebral vascular architecture is exquisitely tailored to functional demands. The spatial organization of arteriovenous networks demonstrates remarkable regional specialization: in high-metabolic regions such as the sensorimotor cortex, dense and tightly coordinated arterial-venous patterning facilitates superior hemodynamic performance, whereas territories with relatively sparse venous networks demonstrate reduced circulatory efficiency and weaker arteriovenous synchrony. This vascular patterning mirrors evolutionary adaptations in rodents, whose sensorimotor systems are critical for whisker-based environmental navigation and complex motor behaviors (Grandjean et al., 2023; Xiong et al., 2017), and have evolved optimized vascular networks to meet their high-energy demands. Collectively, these observations provide compelling hemodynamic evidence for the fundamental principle of structure-function adaptation.

In summary, our analytical framework represents a conceptual advance in cerebrovascular research. Rather than focusing on isolated vascular parameters or static perfusion metrics, our approach captures the dynamic coordination between arterial supply and venous drainage across the entire vascular network. This paradigm shift not only overcomes the limitations of traditional artery-centric analyses but also provides a powerful framework for characterizing the spatiotemporal dynamics of cerebral microcirculation.

### 4.4 Evaluating Microcirculatory Dysfunction Induced by Ischemic Stroke

Ischemic stroke has long been considered a disorder of arterial insufficiency, with therapeutic paradigms centered on vascular recanalization and restoration of cerebral blood flow (Hilkens et al., 2024; Kropf et al., 2025). However, nearly half of patients still experience persistent hypoperfusion, motor dysfunction, and incomplete cognitive recovery despite successful reperfusion therapy, challenging the sufficiency of arterial-centric pathophysiological models. Recent investigations increasingly implicate venous outflow obstruction as a critical contributor to stroke pathogenesis (Li et al., 2023; Wang et al., 2023). The interplay between arterial insufficiency and venous congestion provides a possible framework for explaining the clinical paradox of successful reperfusion but poor functional outcomes. Based on this integrative perspective, we propose that stroke-induced ischemia involves microcirculatory failure, mediated by dysfunction of the arteriovenous circulation. To validate this hypothesis, we developed a novel assessment framework quantifying arteriovenous fluctuation coupling and implemented it in an acute ischemic stroke model to characterize microcirculatory dynamics.

Our findings demonstrate that stroke-induced microcirculatory impairment manifests across both temporal and spatial dimensions. Temporally, post-stroke arteriovenous dynamics exhibited delayed state transitions, consistent with previous reports describing cerebrovascular sluggishness and impaired vascular responsiveness following ischemic injury (Østergaard et al., 2016; Wang et al., 2023). Spatial analyses revealed marked regional heterogeneity in ischemic vulnerability: visual cortical regions characterized by relatively sparse arteriovenous integration exhibited more severe disruption of circulation dynamics, whereas somatosensory regions with denser mixed vascular architecture exhibited comparatively preserved function. Although these findings do not establish causality, they suggest that regions with stronger arteriovenous integration may possess greater hemodynamic resilience, potentially through enhanced collateral compensation or improved redistribution of blood flow following ischemia (Glück et al., 2024; Xiong et al., 2017; Xue et al., 2025). Quantitative analyses further demonstrated that artery-dominant vascular composition correlated positively with ischemic lesion extent, whereas higher proportions of mixed arteriovenous regions correlated inversely with lesion severity and positively with the penumbra/core ratio. These findings should be interpreted cautiously, as they likely reflect region-specific vascular organization rather than direct causal protection.

Of note, the microcirculation-based method yielded ischemic lesion maps highly consistent with those from the conventional CBF approach, confirming its comparable performance in identifying ischemic lesions. Beyond standard CBF, our microcirculation framework further characterizes ischemic injury by integrating temporal dynamic patterns and spatial heterogeneity, thereby providing multidimensional diagnostic insights unavailable from conventional perfusion analysis. Furthermore, our approach achieves a more pronounced separation between the ischemic penumbra and infarct core, facilitating a more rigorous assessment of ischemic heterogeneity. Collectively, the proposed strategy not only preserves the lesion-detection capability of standard CBF approaches but also expands the analytical dimensions for evaluating stroke-induced injury, offering a novel perspective for the comprehensive diagnosis of acute stroke.

### 4.5 Limitations

Several limitations should be acknowledged. First, the present framework is derived primarily from cortical surface laser speckle imaging and therefore does not directly capture deep-brain or capillary-level microcirculatory dynamics. Accordingly, the term “microcirculation” in this study should be interpreted as large-scale cortical microvascular coordination rather than direct visualization of capillary exchange. Second, the arteriovenous circulation index represents an indirect surrogate measure of vascular synchronization rather than a direct quantification of perfusion efficiency or oxygen delivery. Third, because all imaging experiments were performed under isoflurane anesthesia, anesthesia-related modulation of vascular oscillations and neurovascular coupling may influence the observed dynamics. Fourth, although signal purification analyses demonstrated strong robustness, definitive validation against ground-truth vascular labeling methods, such as fluorescent angiography or two-photon microscopy, was not performed. Finally, the observed associations between vascular architecture and ischemic vulnerability remain correlational and should not be interpreted as evidence of direct causal protection.

### 4.6 Conclusion

Using a co-fluctuation analysis framework, we developed a dynamic arteriovenous circulation index that quantifies the temporal coordination between arterial inflow and venous outflow during cerebral blood flow activity. Application of this framework in ischemic stroke mice demonstrated that disruption of arteriovenous synchronization is associated with altered microcirculatory dynamics and regional ischemic vulnerability. These findings provide a systems-level perspective on cerebrovascular dysfunction that extends beyond conventional artery-centric perfusion analyses and may offer a useful framework for future investigation of microvascular pathophysiology in ischemic stroke.

## Credit authorship contribution statement

Conceptualization Bochao Niu, Hongyan Gong, Data Collection and Curation Bochao Niu, Hongyan Gong, Yang Yuan, Methodology Bochao Niu, Writing–Original Draft Bochao Niu, Writing-Review Benjamin Klugah-Brown, Quandan Tan, Guoliang Zhu, Yapeng Lin, Junli Hao, Kejie Chen, Lingling Wang, Zhe Kang Law, Supervision Jie Yang, Funding acquisition Jie Yang, Hongyan Gong, Yanlin Bi. All authors read and approved the submitted version of the manuscript.

## Code Availability

To access the broader scientific exploration, the customized MATLAB scripts mentioned in this research are available in Github (https://github.com/JoeySmartWei/Microcirculation-Assessment-based-on-Arteriovenous-Co-fluctuation-Analysis). The manual vascular segmentation datasets and the resting-state datasets of healthy rodents are publicly available in the Science Data Bank (ScienceDB) at https://www.scidb.cn/anonymous/QmZhMjZq. The MCAO stroke datasets are restricted due to privacy concerns, but can be obtained from the corresponding author upon reasonable request.

## Declarations

The author(s) declared no potential conflicts of interest with respect to the research, authorship, and/or publication of this article.

## Acknowledgments

This work was supported by the National Natural Science Foundation of China (NSFC82571515), the National Key Research and Development Program of China (2026YFE0215600), the Sichuan Provincial Science and Technology Department (2023YFS0042), the Health Commission of Sichuan Provincial (24LCYJZD04), and the Qingdao Science and Technology Program for Benefiting People Special Project (No.25-1-5-smjk-16-nsh).

## Notes

https://github.com/JoeySmartWei/Microcirculation-Assessment-based-on-Arteriovenous-Co-fluctuation-Analysis

https://www.scidb.cn/anonymous/QmZhMjZq

### Competing Interest Statement

The authors have declared no competing interest.

