## Supplementary Tables and Supplementary Figures for "Assessing Microcirculation Impairment in Ischemic Stroke Mice Using Arteriovenous Co-fluctuation Analysis"

**Bochao Niu^1 #^, Yanlin Bi^2,3#^,** [**Benjamin Klugah-Brown**](https://pubmed.ncbi.nlm.nih.gov/?term=Klugah-Brown+B&cauthor_id=37931478)**^4,5^, Yang Yuan^2^, Quandan Tan^1^, Guoliang Zhu^1^, Yapeng Lin^6^, Junli Hao^7^, Kejie Chen^8^, Lingling Wang^9^, Zhe Kang Law^10,11^, Hongyan Gong^2,3^*, Jie Yang^1^***

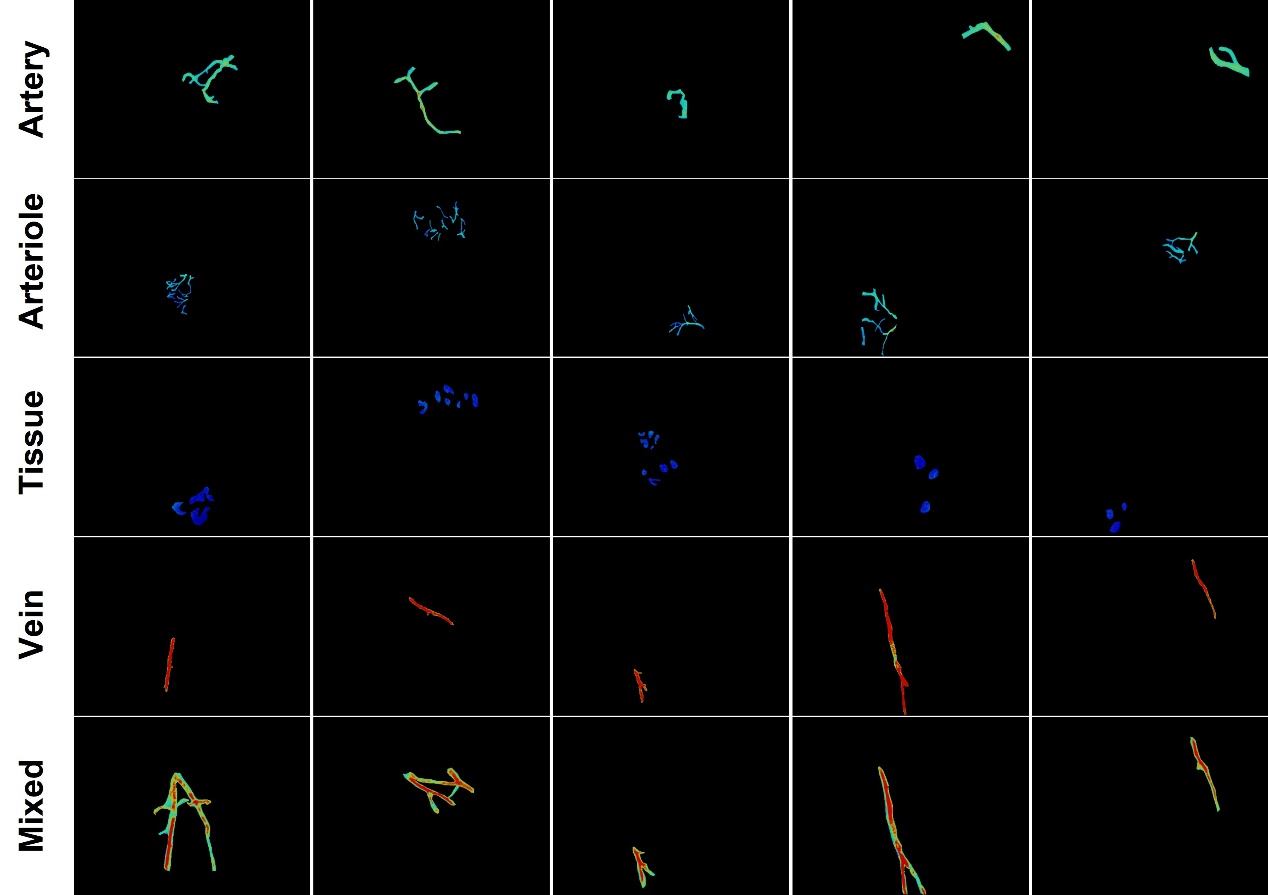

**Fig. S1** Manually segmented masks of different vessel types.

**Table S1.** **Statistics of segmented vascular counts in the sensorimotor cortex.**

| Num.  Vessels | R1 | R2 | R3 | R4 | R5 | R6 | R7 | R8 | R9 | R10 | R11 | R12 | R13 | R14 | R15 | R16 | R17 | R18 | R19 |
| --- | --- | --- | --- | --- | --- | --- | --- | --- | --- | --- | --- | --- | --- | --- | --- | --- | --- | --- | --- |
| Vein | 3 | 2 | 4 | 2 | 1 | 2 | 2 | 3 | 1 | 3 | 3 | 3 | 2 | 2 | 2 | 2 | 3 | 1 | 3 |
| Artery-vein | 1 | 2 | 4 | 3 | 1 | 2 | 2 | 2 | 3 | 2 | 3 | 3 | 4 | 3 | 3 | 2 | 3 | 3 | 3 |
| Artery | 1 | 4 | 4 | 2 | 2 | 3 | 4 | 2 | 0 | 0 | 3 | 2 | 2 | 3 | 1 | 4 | 2 | 3 | 4 |
| Arteriole | 2 | 3 | 2 | 2 | 1 | 3 | 1 | 1 | 1 | 3 | 1 | 3 | 1 | 2 | 2 | 3 | 2 | 2 | 2 |
| Tissue | 1 | 3 | 2 | 2 | 1 | 3 | 2 | 1 | 1 | 3 | 3 | 3 | 3 | 3 | 2 | 2 | 3 | 3 | 3 |

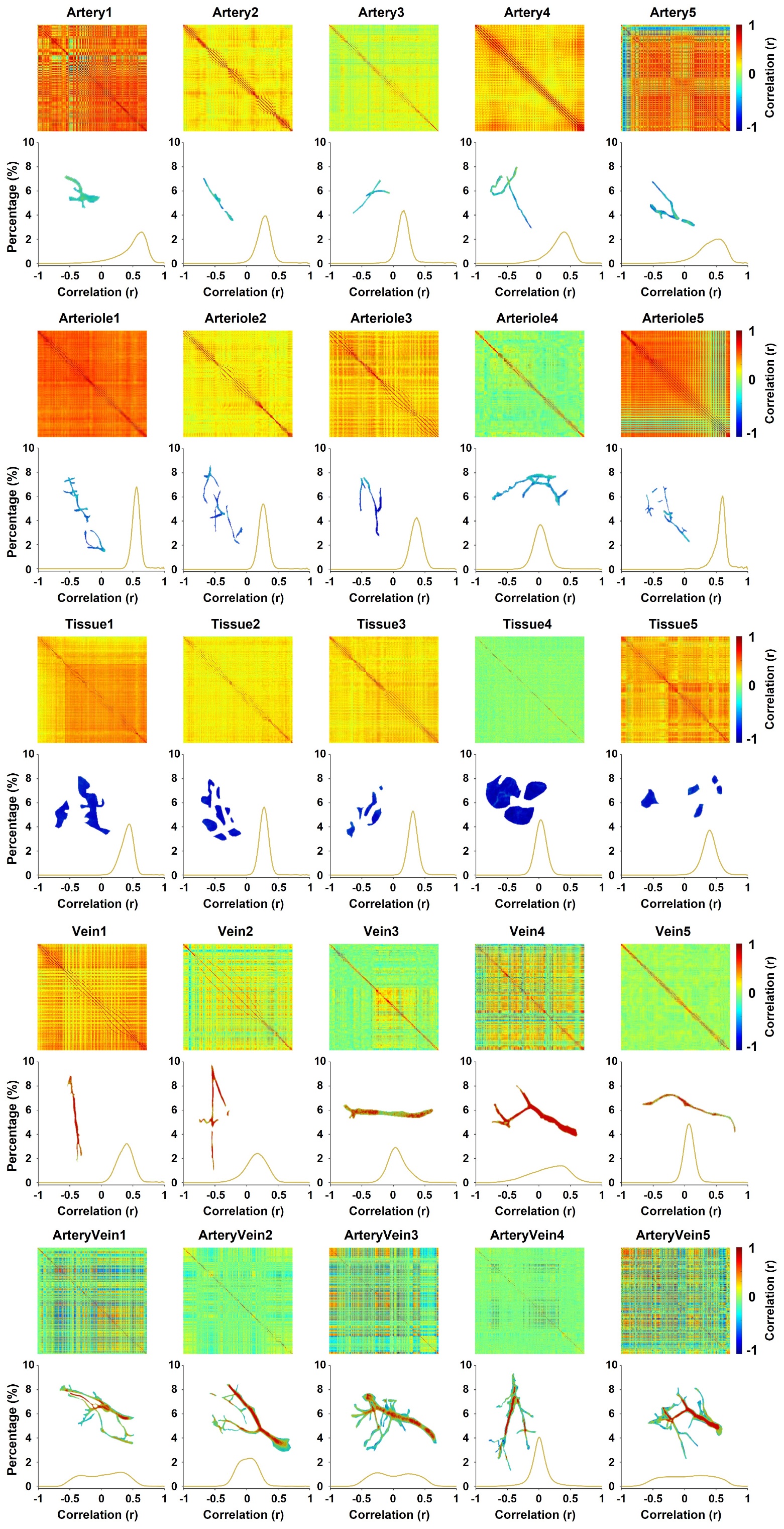

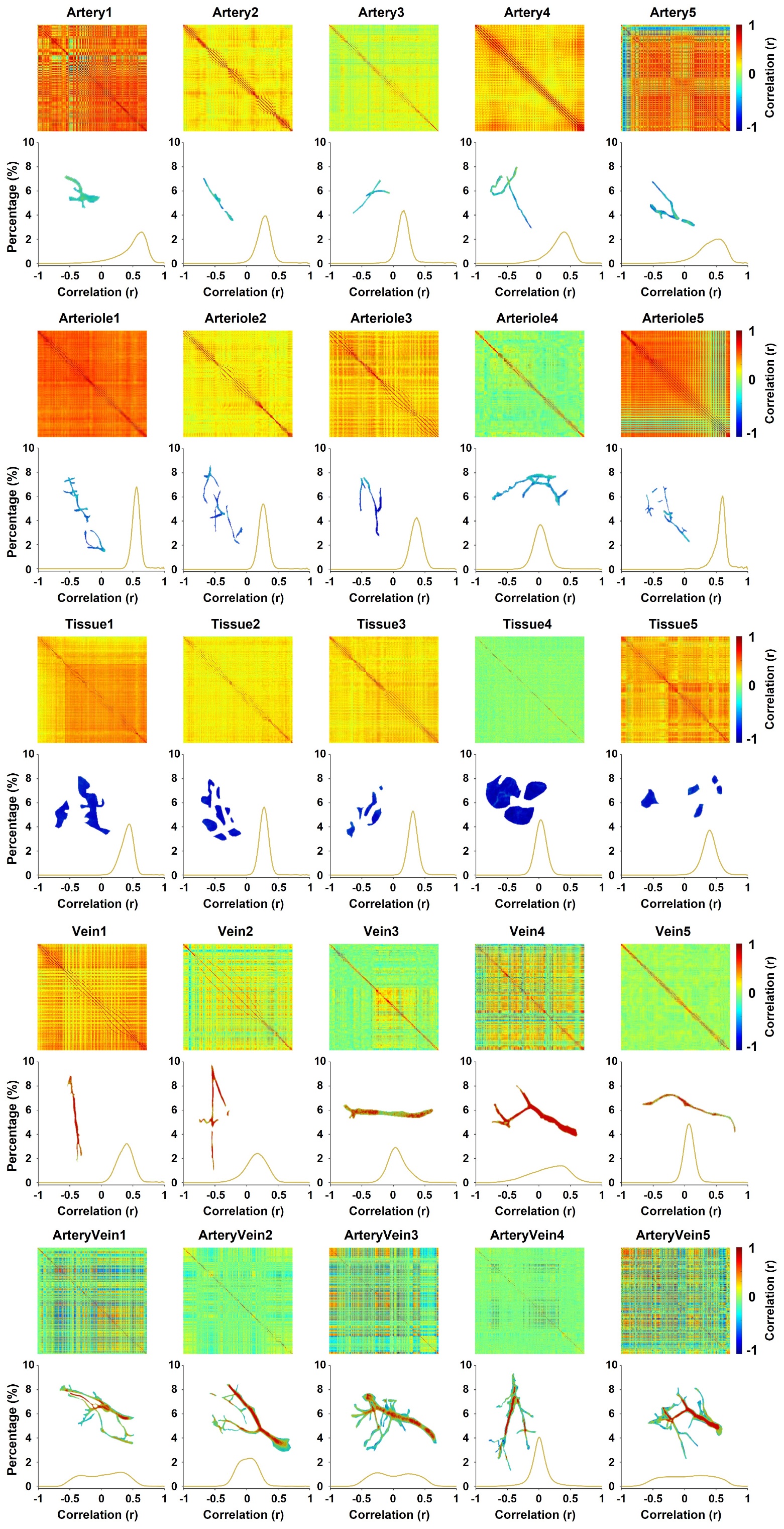

**Fig. S2** Correlation matrices of whole pixel signals within different vascular masks. Arterioles and cerebral tissue masks exhibit the highest internal homogeneity in pixel correlation matrices, large arteries are intermediate, and veins and arteriovenous mixture masks show the lowest internal homogeneity.

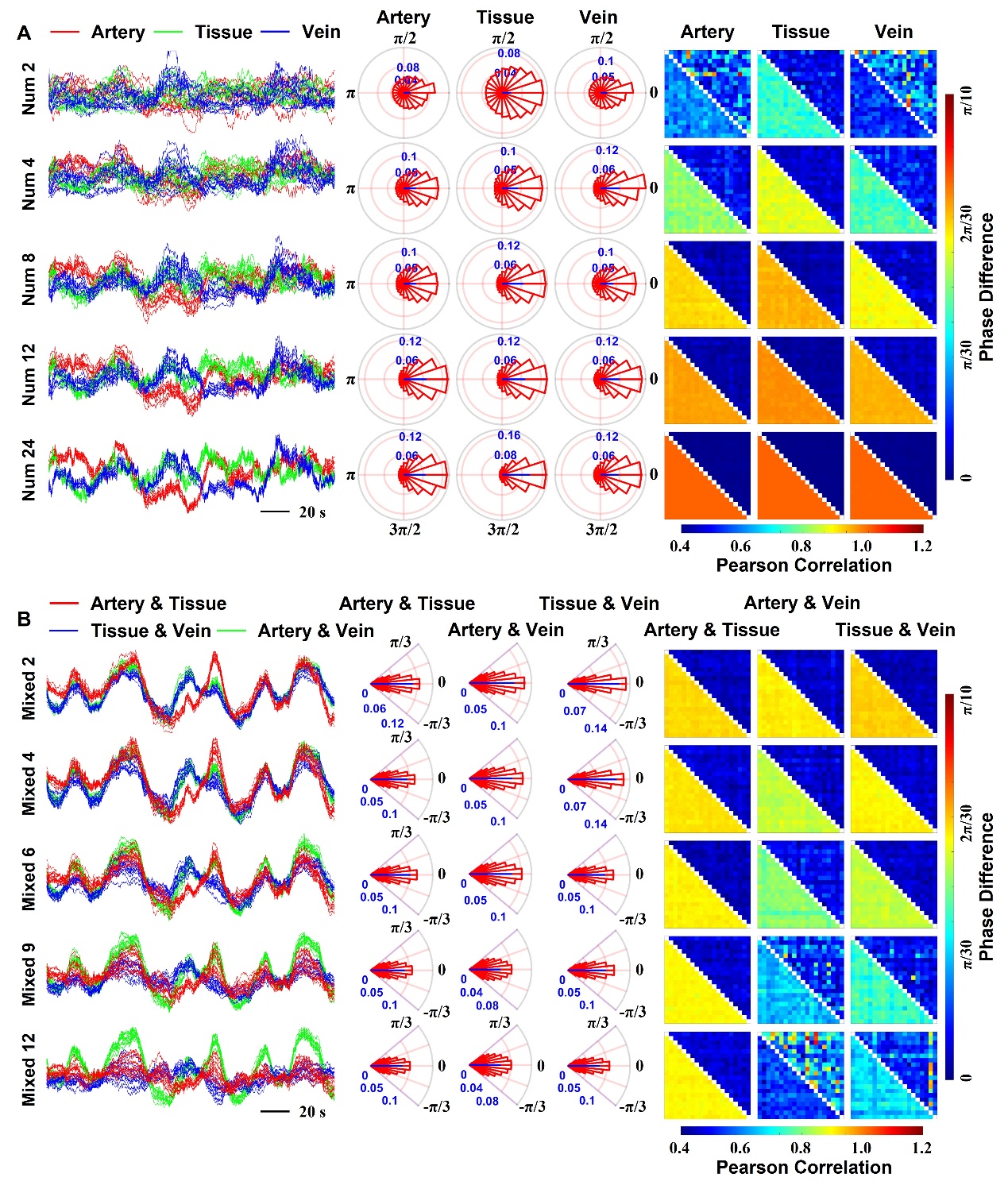

**Fig. S3 A.** When ROI signals are purified using a small set of pixels, phase differences are widely scattered. As the pixel count increases, these differences converge to 0°, and the Pearson correlation approaches 1. **B.** In arterial–tissue mixtures containing 2–12 mixed pixels, phase differences remain stable near 0°. However, for arterial–venous mixtures, once the number of mixed pixels exceeds 6, phase differences diverge significantly from 0°, accompanied by a rapid drop in Pearson correlation to below 0.8.

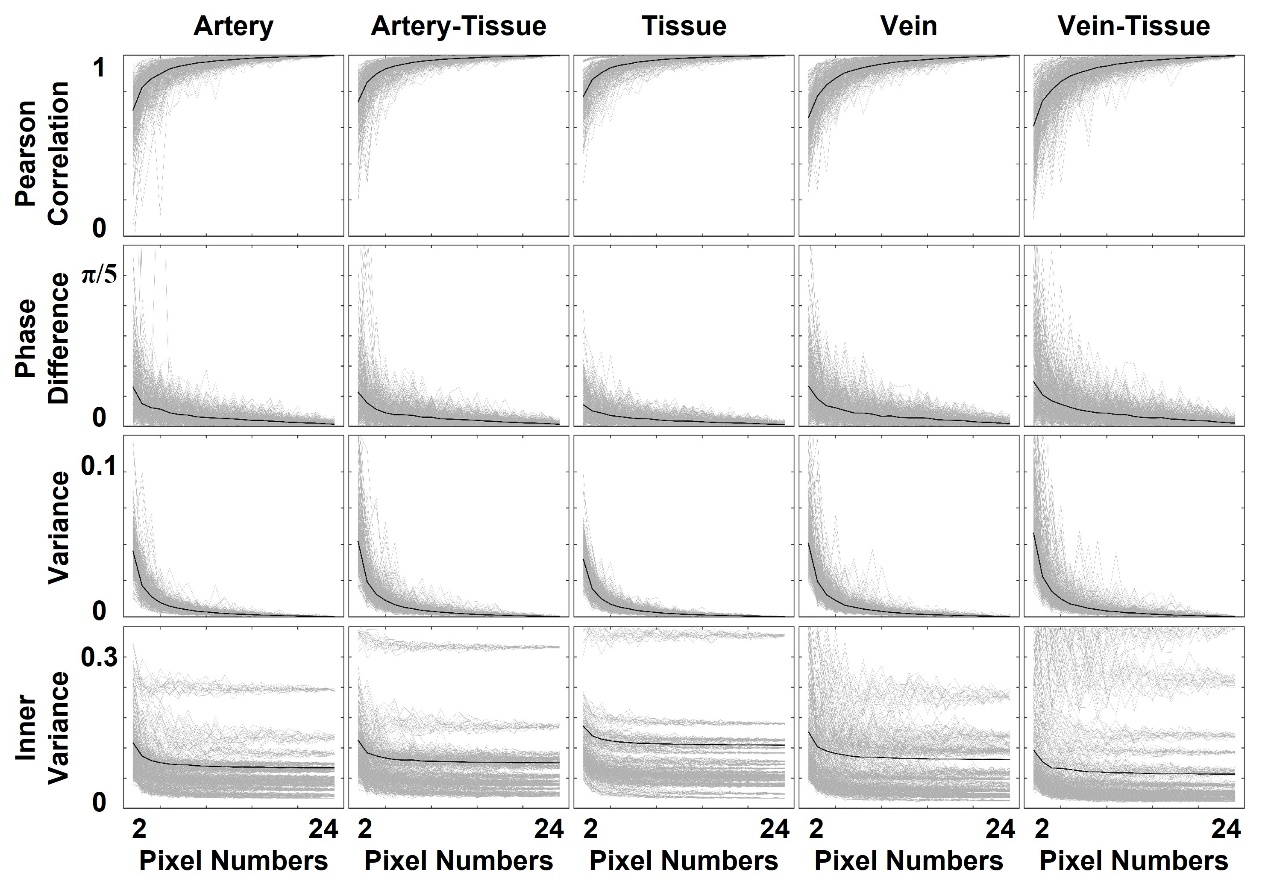

**Fig. S4** Purification performance across varying pixel counts. As the number of randomly selected pixels from individual ROIs increases, the purified signals converge rapidly toward the reference: the correlation coefficient toward 1, the phase difference toward 0°, coinciding with a progressive reduction in both inter-signal variance and intrinsic signal variability to their minima.

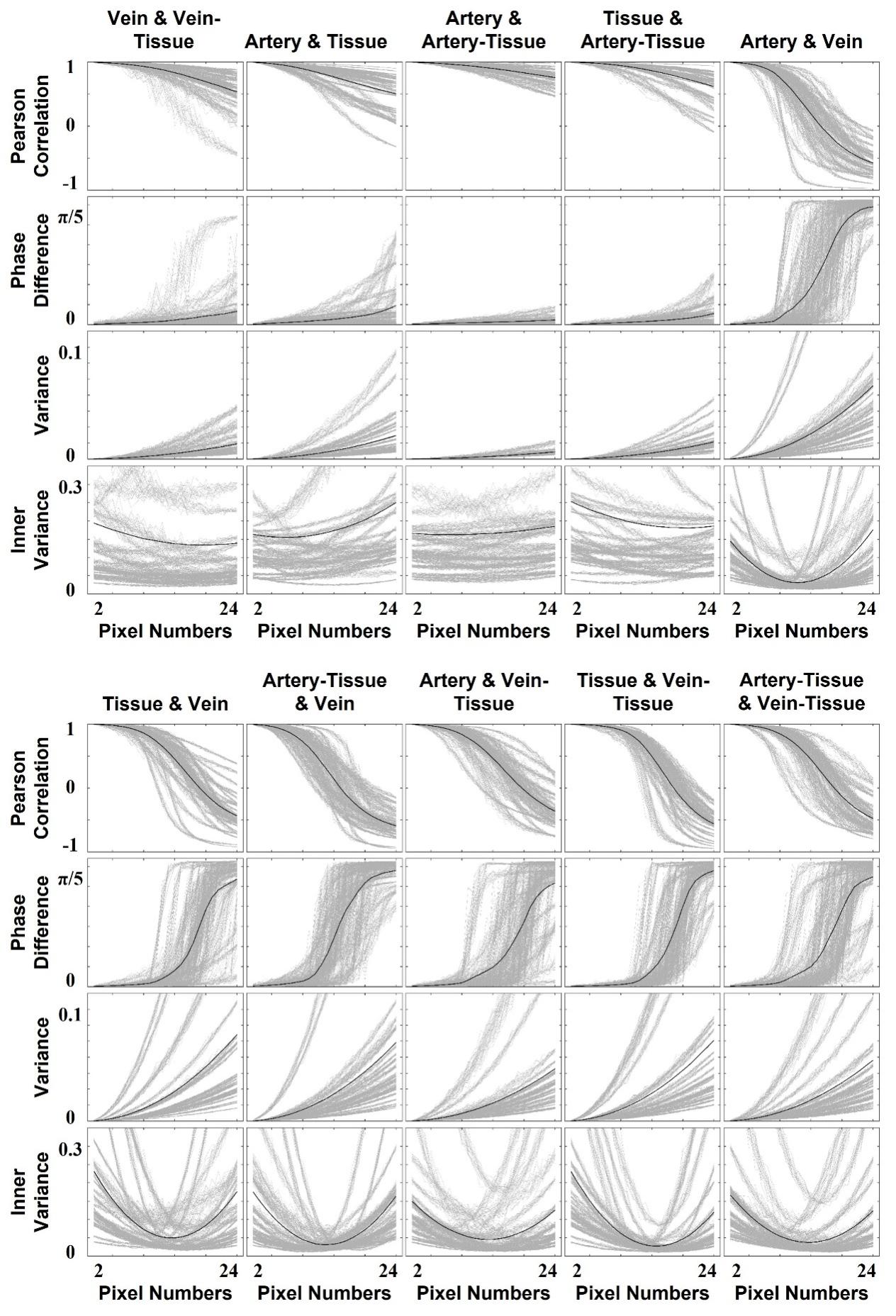

**Fig. S5** Mixing process for different vascular ROI signals. When homogeneous cerebral tissue or artery–tissue signals are mixed into arterial ROIs, the signal characteristics remain stable relative to the reference across increasing levels of pixel mixing. In contrast, as the fraction of heterogeneous venous or vein–tissue pixels rises to 50%, both the Pearson correlation and intrinsic signal variability reach to their minima, while the phase difference approaches a critical transition point. Beyond this threshold, further increases in heterogeneous pixels drive the Pearson correlation into strong negative values, cause the phase difference to rapidly converge to 180°, and restore intrinsic signal variability.

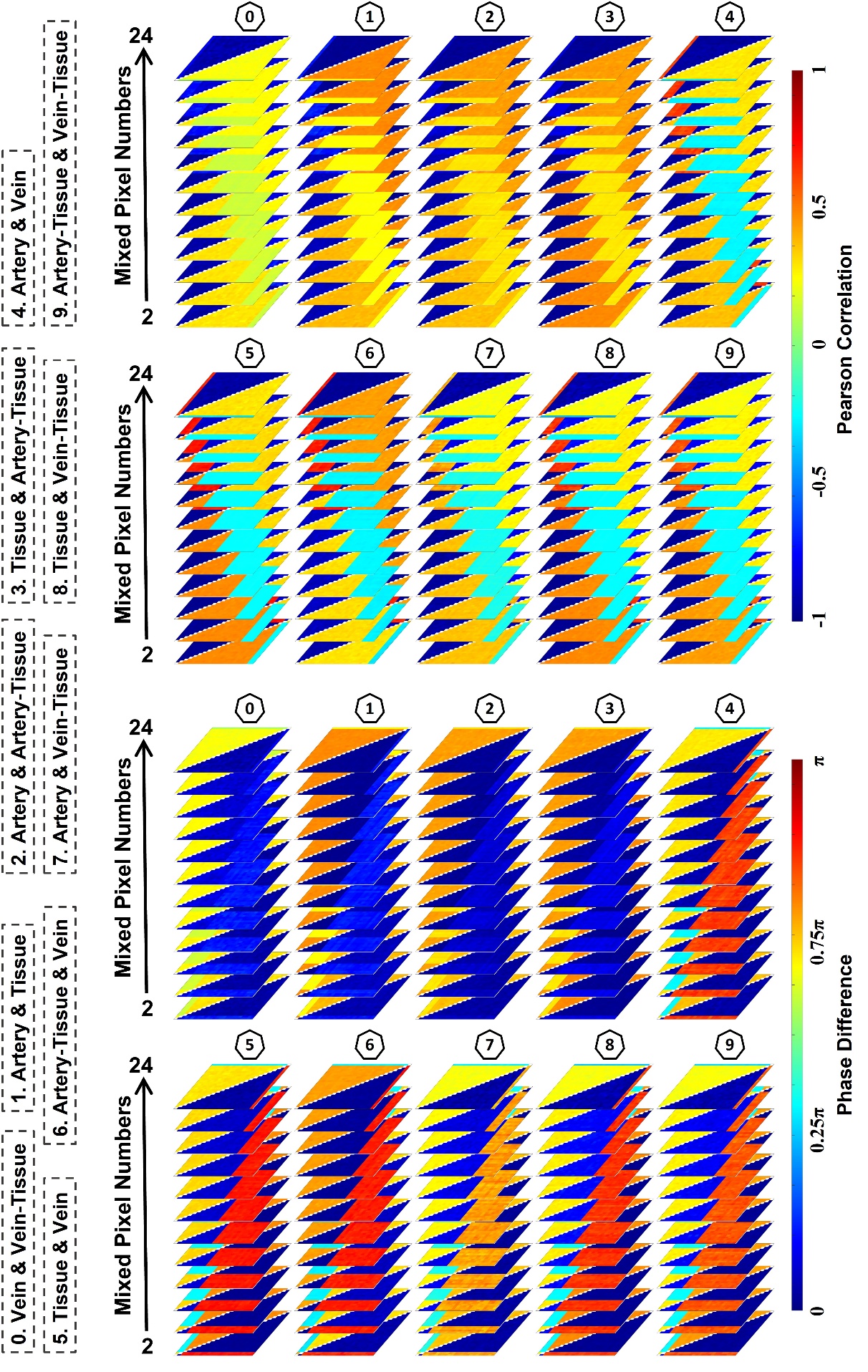

**Fig. S6** Matrices of Pearson correlation and phase difference for distinct vascular ROI mixtures.

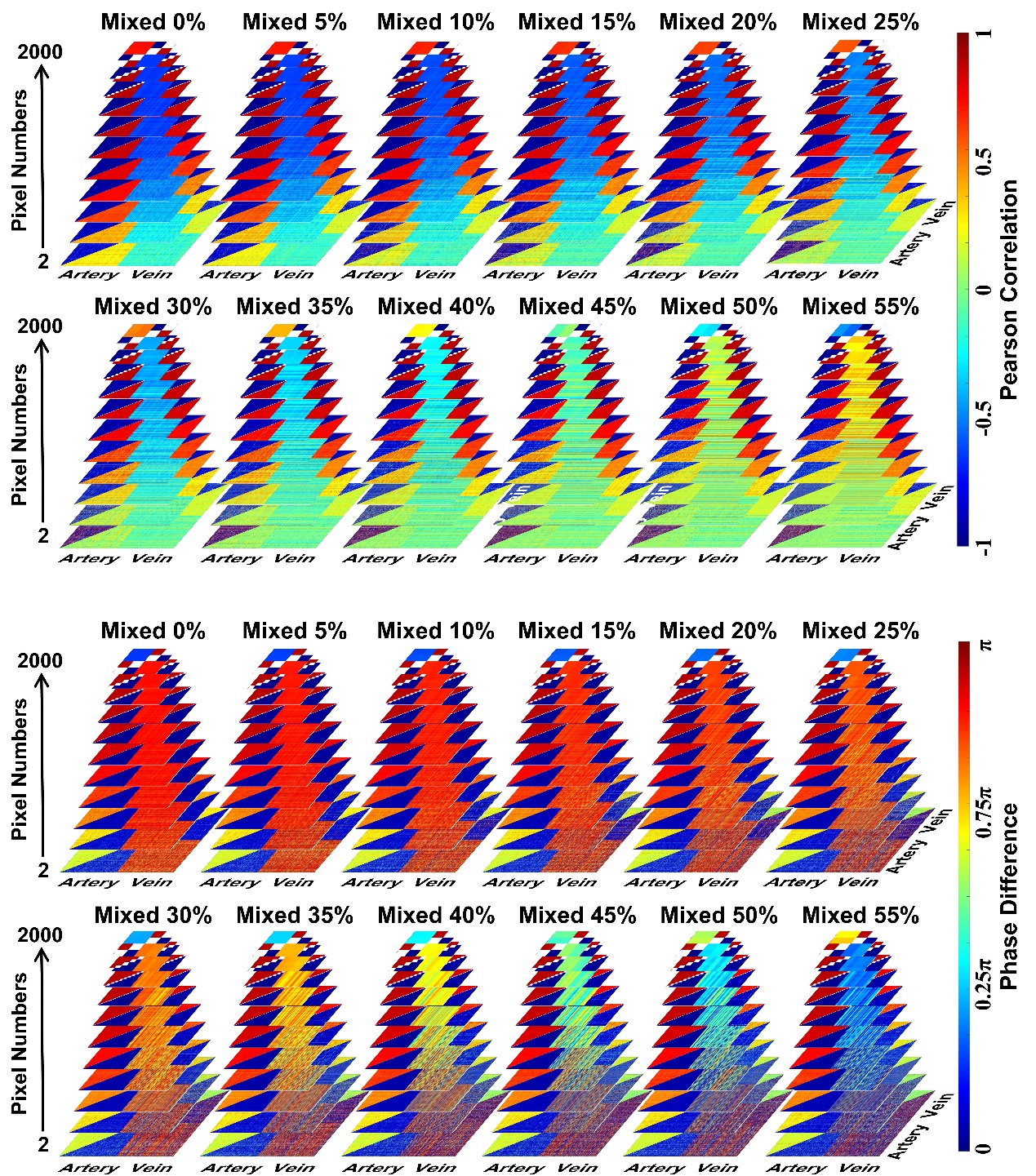

**Fig. S7** Purification efficacy of arteriovenous mixed signals across varying mixing ratios. The method demonstrates robust performance at mixing ratios below 20%, yielding a strong negative Pearson correlation below -0.9 and a phase difference converging to 180°. However, at ratios exceeding 40%, purification performance significantly degrades: the correlation weakens to above -0.3, and the phase difference deviates substantially, remaining below 135°.

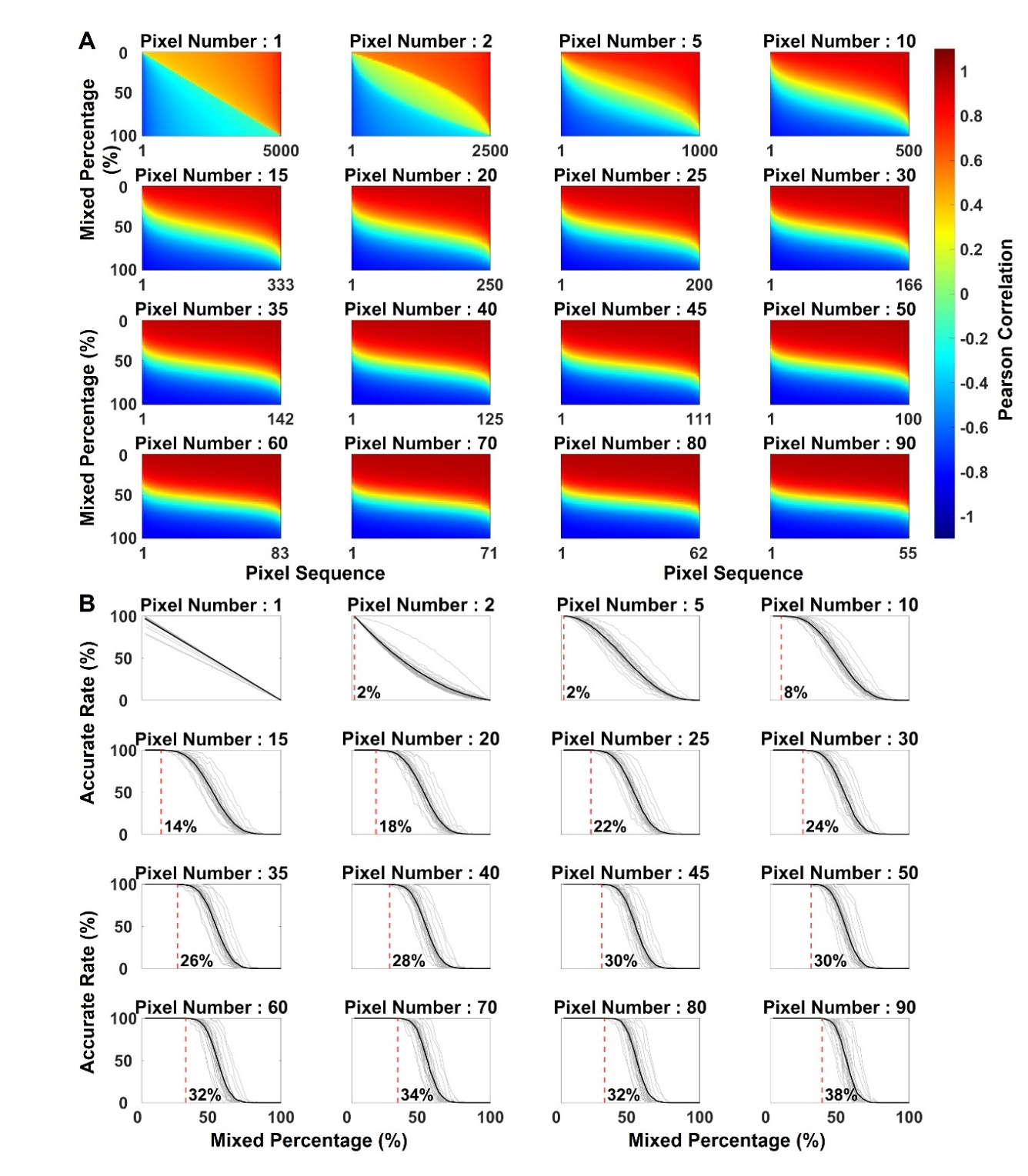

**Fig. S8** Quantitative analysis of the impact of arteriovenous mixing ratio on signal purification. **A.** Increasing the arteriovenous mixing ratio (0-100%) degrades purification performance and reduces the fraction of strongly correlated signals. Conversely, increasing the number of purification pixels enhances performance and increases the fraction of strongly correlated pixels. **B.** Noise tolerance, measured at the 90% strong correlation threshold, increases with the number of purification pixels.

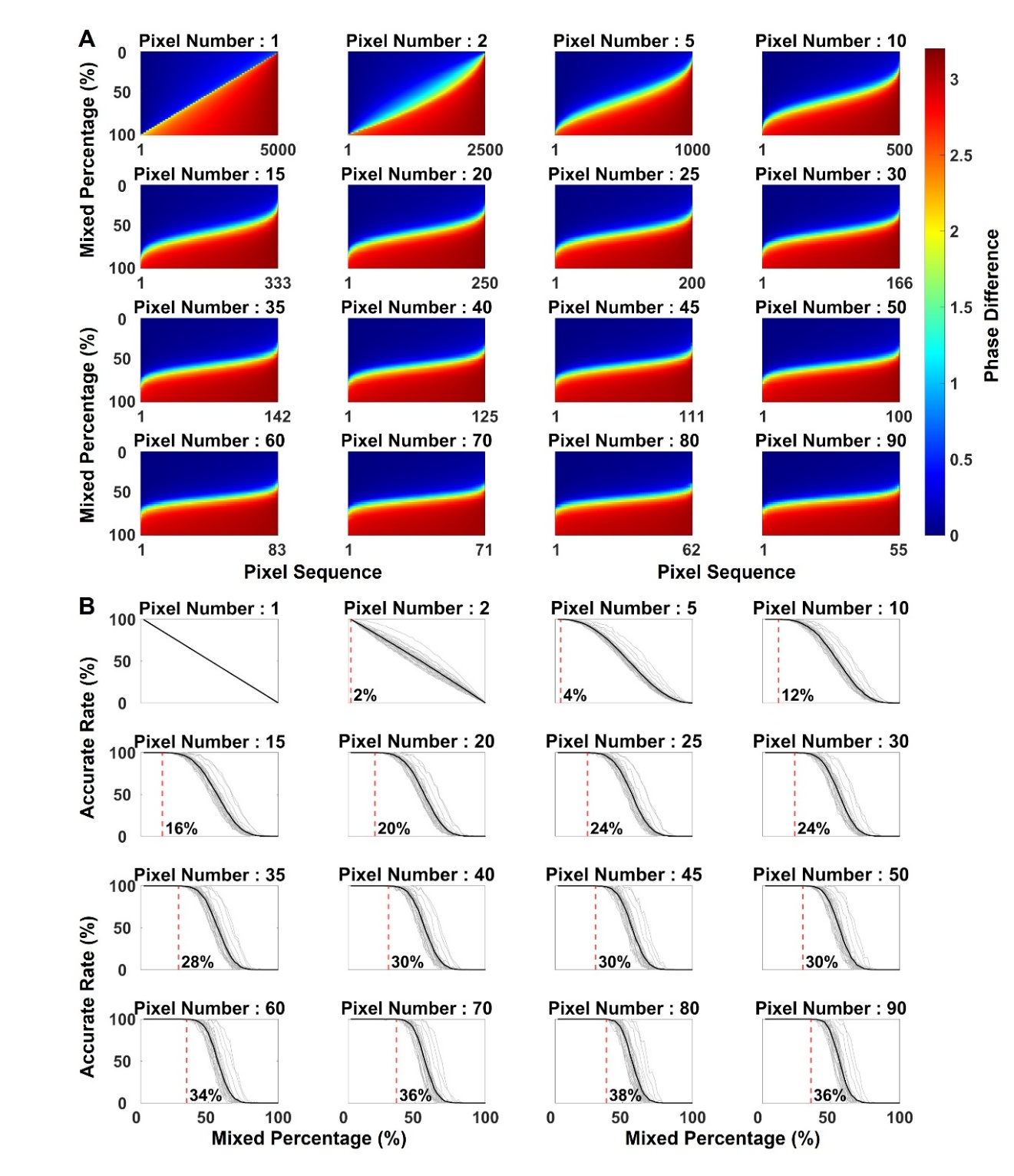

**Fig. S9** Quantitative analysis of arteriovenous mixing effects on signal purification. **A.** Increasing the mixing ratio (0–100%) reduces purification efficacy and decreases the fraction of low phase differences between mixed and reference signals. In contrast, increasing the number of purification pixels improves performance and boosts the fraction of low-phase-difference signals. **B.** The noise tolerance threshold, defined as the noise level at which the fraction of low phase difference drops to 90% of baseline, increases with the number of purification pixels.

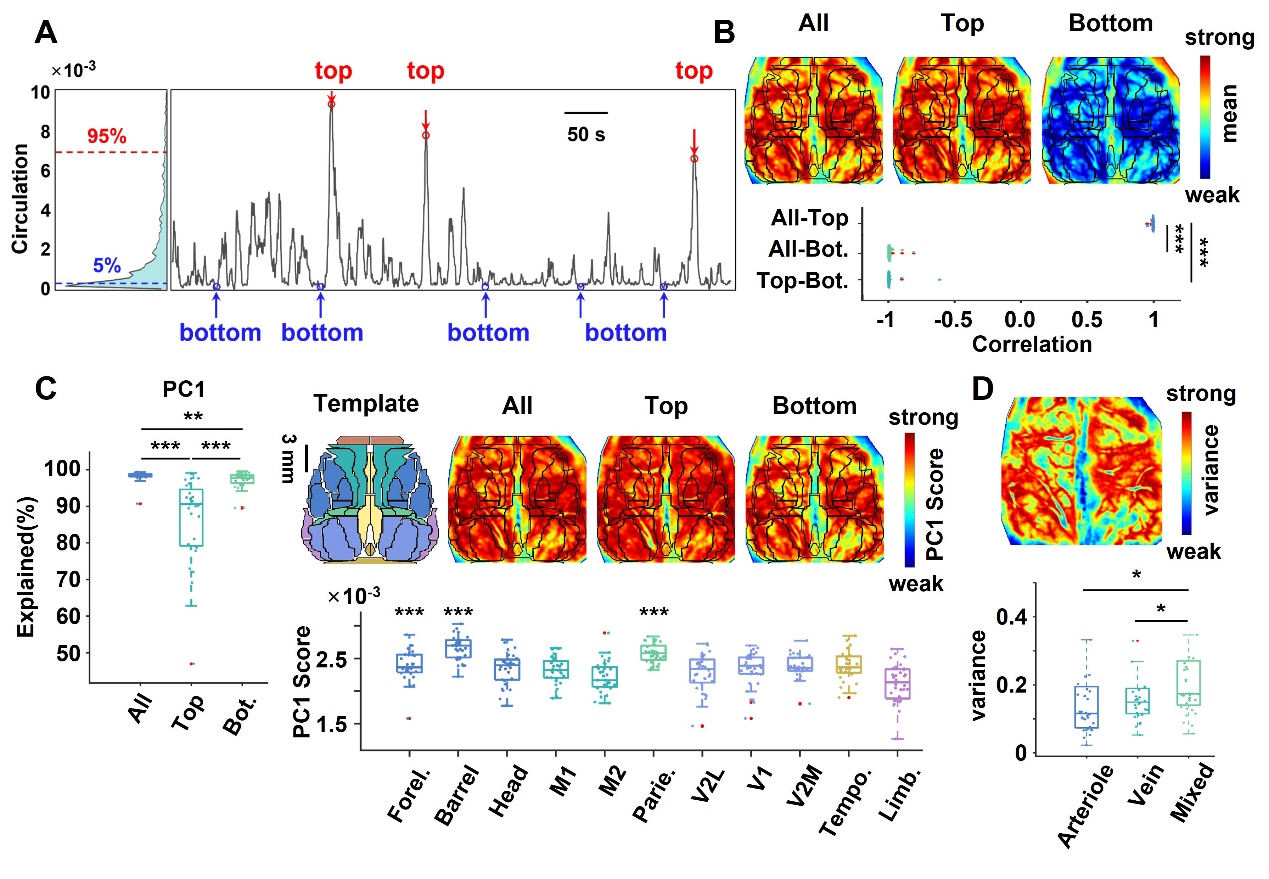

**Fig. S10** Spatiotemporal characteristics of whole-cerebral blood circulation activity in rats. **A, B.** The time series of arteriovenous fluctuation coupling exhibits a right-skewed distribution. Timepoints corresponding to the top 5% (peaks) and bottom 5% (troughs) of this distribution were extracted for further analysis. The cortical activity pattern at peaks closely resembles that of the full time series, but is negatively correlated with that at troughs. **C.** The spatial pattern of cortical circulation activity is highly consistent across peak, trough, and full signals, characterized by the strongest activity in the sensorimotor cortex and the weakest in the temporal and limbic cortices. Interregional parameter differences are examined by one-way ANOVA (n = 30), followed by pairwise t-tests with FDR correction using V2L as the reference. **D.** Arteriovenous mixed regions exhibited significantly higher temporal variability than arteriole and vein compartments. Comparisons of individual items within timepoints and vascular components are assessed via paired t-test with FDR adjustment (n = 30). *p < 0.05, **p < 0.01, ***p < 0.001 in B, C, and D, *** indicates p < 0.001 whose PC1 scores are greater than secondary visual cortex in C. Bot.: bottom

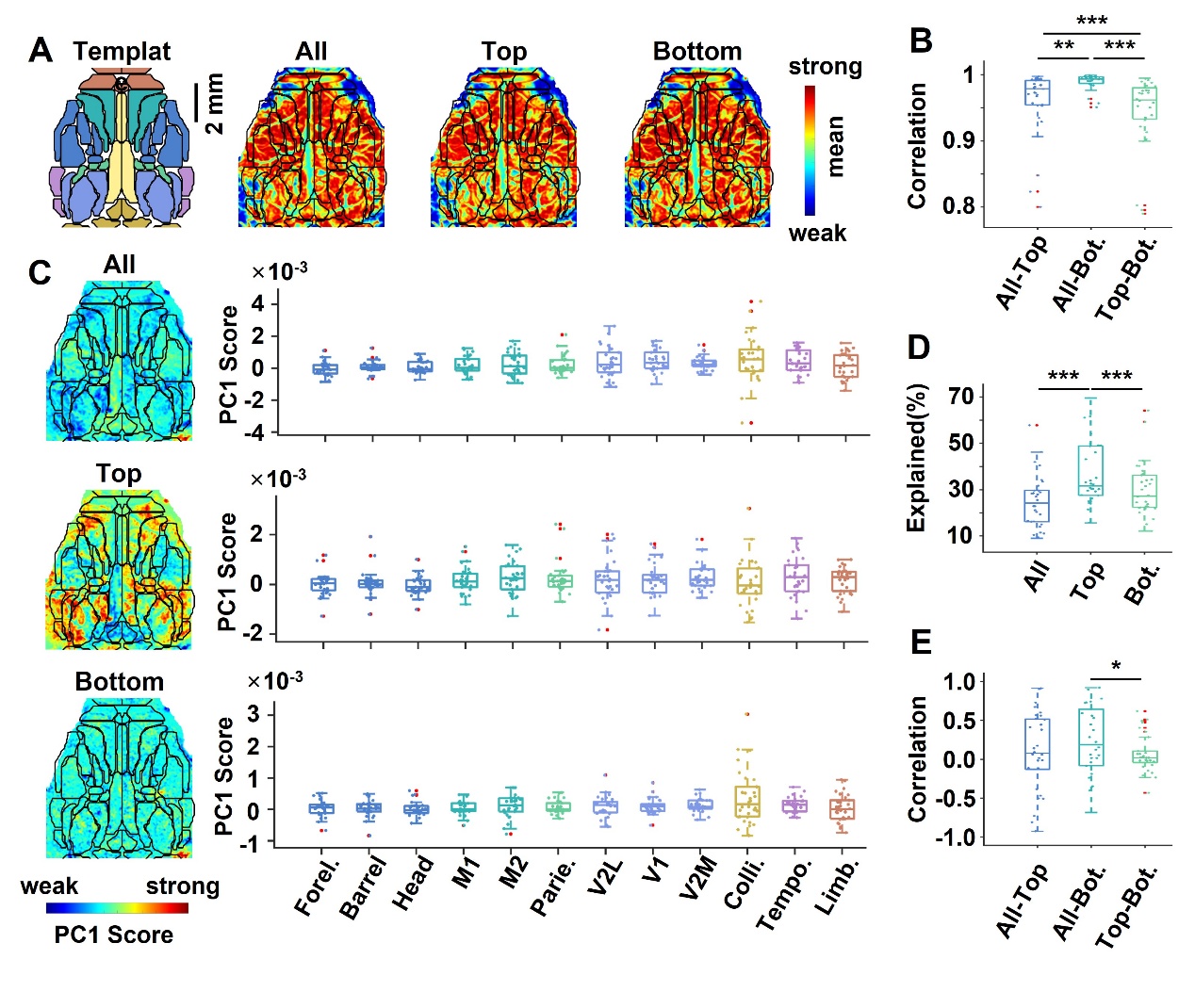

**Fig. S11** Spatial patterns of mouse cortical blood flow activity. **A, B.** Mean blood flow at peaks and troughs correlates strongly with the full signal (r > 0.9). **C–E.** No specific spatial patterns are observed for peaks, troughs, or all timepoints, and weak inter-condition correlations are observed. Interregional parameter differences are examined by one-way ANOVA (n = 30), followed by pairwise t-tests with FDR correction using V2L as the reference. Differences in different timepoints are evaluated via paired t-test with FDR adjustment (n = 30). *p < 0.05, **p < 0.01, ***p < 0.001. Bot.: bottom

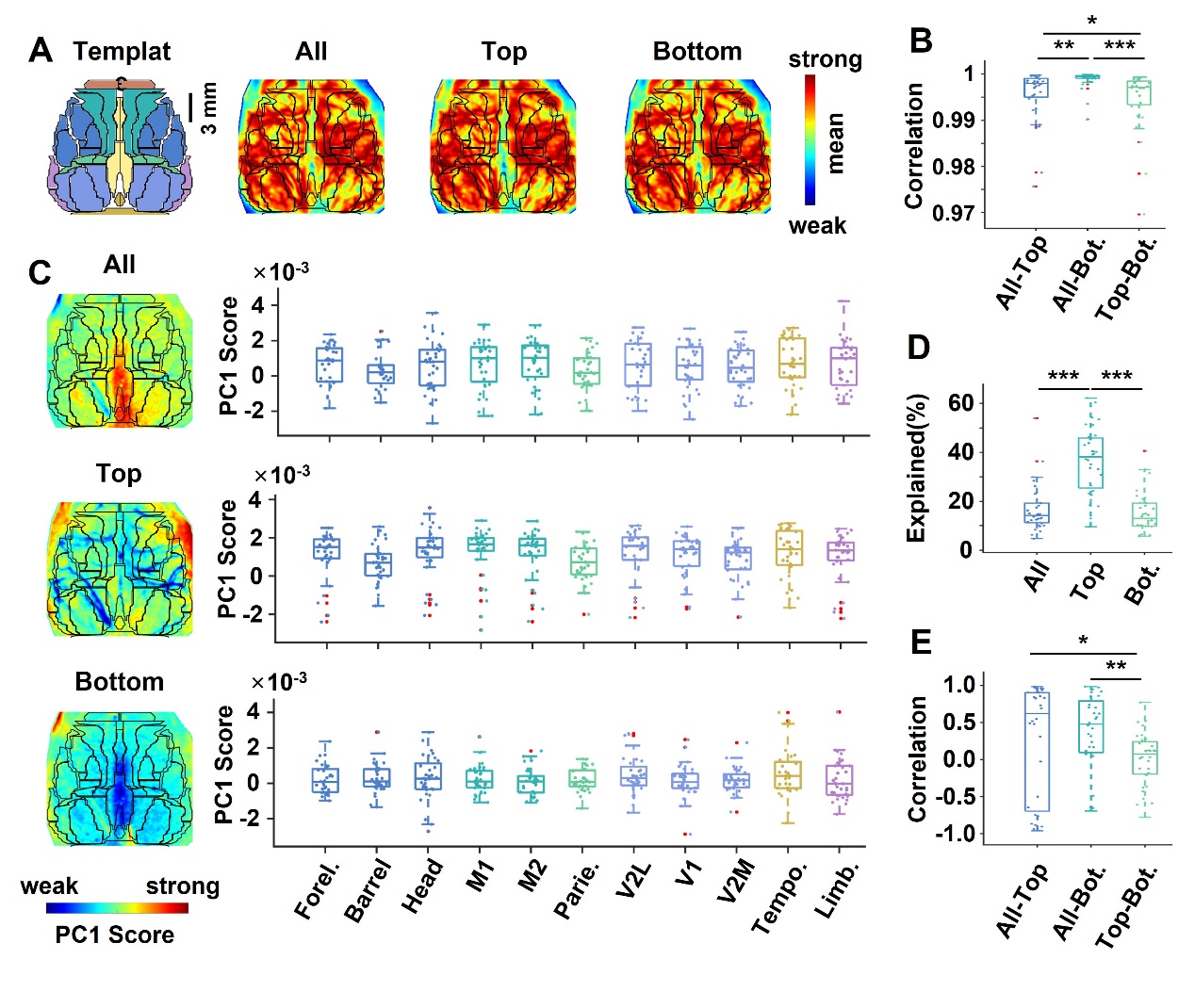

**Fig. S12** Spatial organization of blood flow activity across the entire rat cortex. **A, B.** The mean blood flow topographies at peaks and troughs closely resemble the full signal pattern (r > 0.9). **C–E.** In contrast, peaks, troughs, and the full time series exhibit no distinct spatial structure, showing weak correlations among these temporal states. Interregional parameter differences are examined by one-way ANOVA (n = 30), followed by pairwise t-tests with FDR correction using V2L as the reference. Differences in different timepoints are evaluated via paired t-test with FDR adjustment (n = 30). *p < 0.05, **p < 0.01, ***p < 0.001. Bot.: bottom

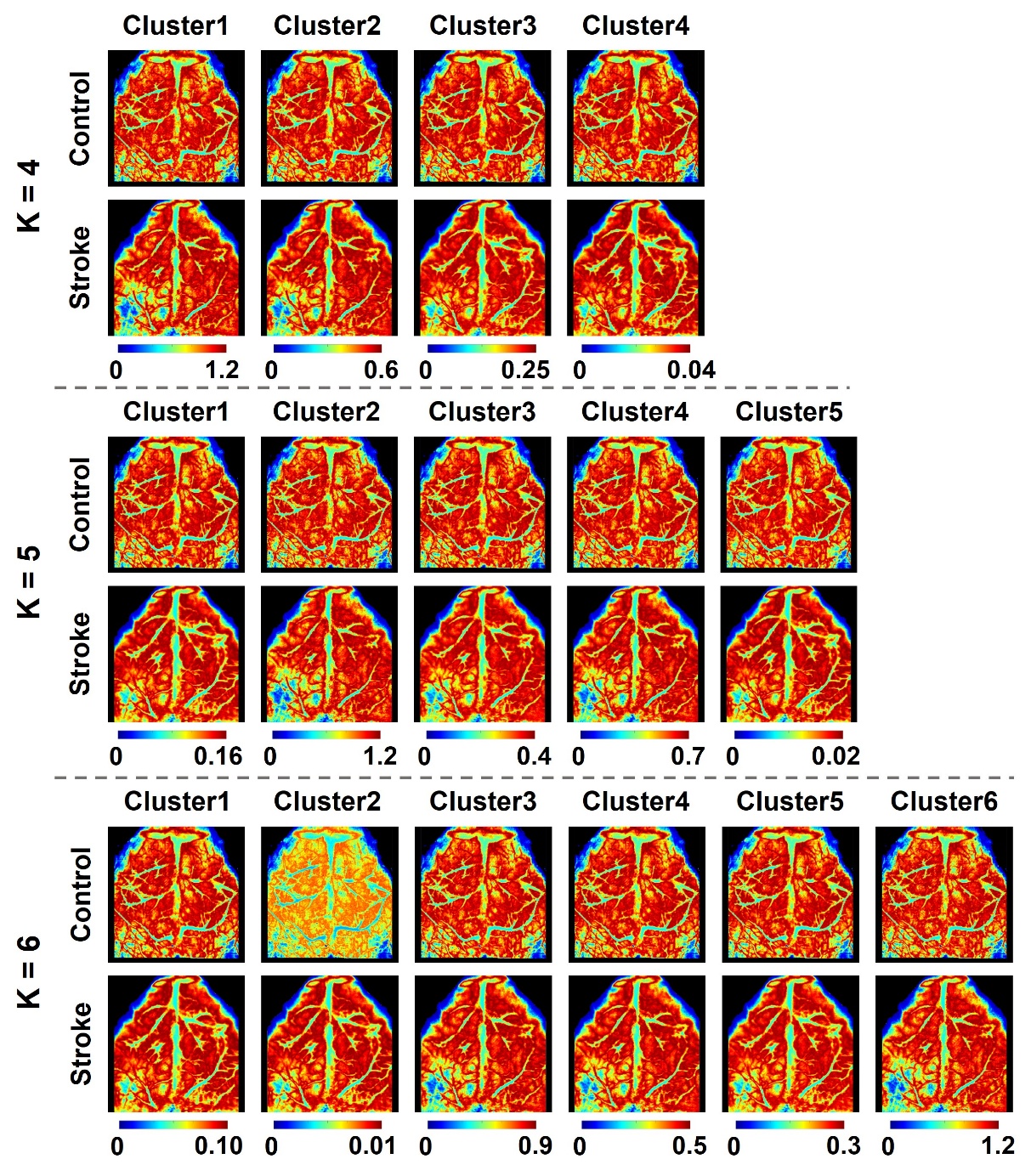

**Fig. S13** Spatial stability of microcirculatory functional states. Whole-brain microcirculatory patterns remain highly consistent across different clustering configurations in both control and stroke groups, showing minimal spatial variation between distinct functional states.

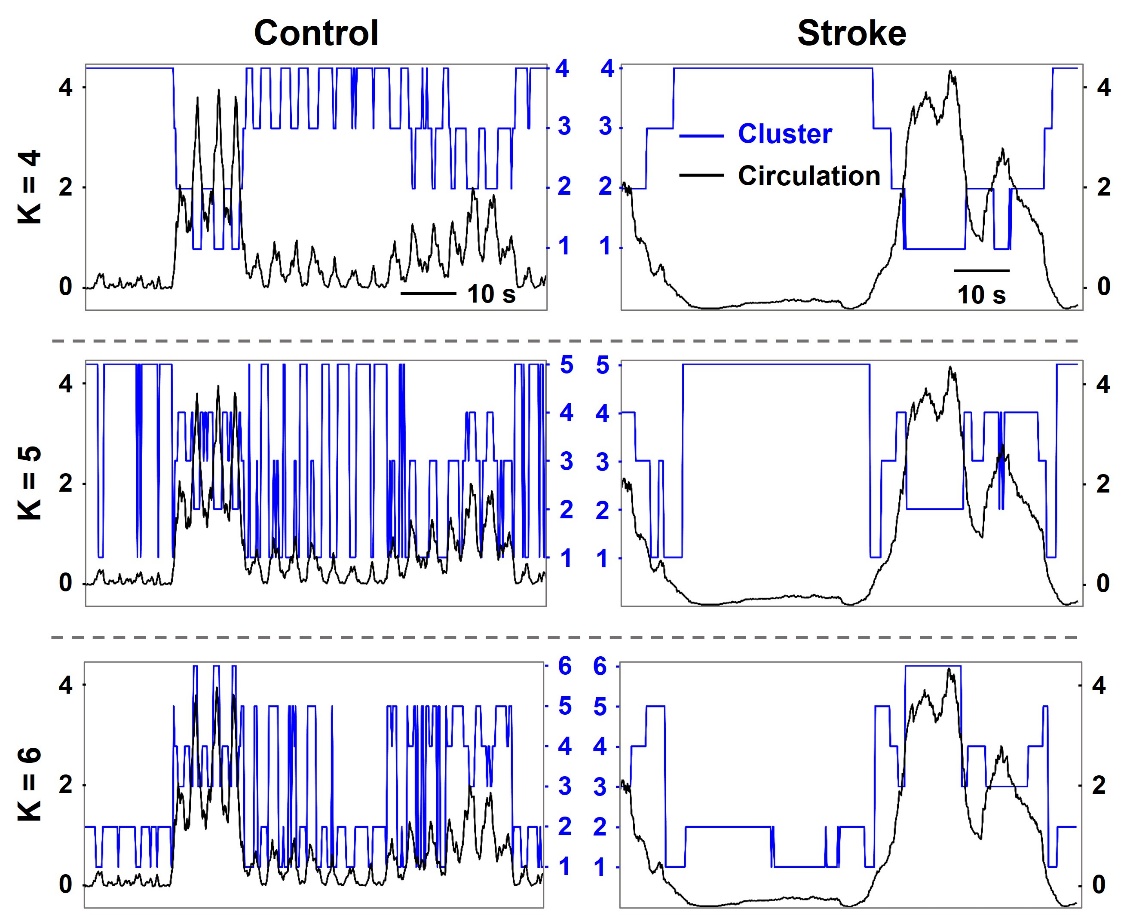

**Fig. S14** Temporal changes of state clustering in microcirculatory activity. Clustering analysis across different numbers of states reveals that stroke prolongs the duration of individual functional states while reducing transition times between distinct states.

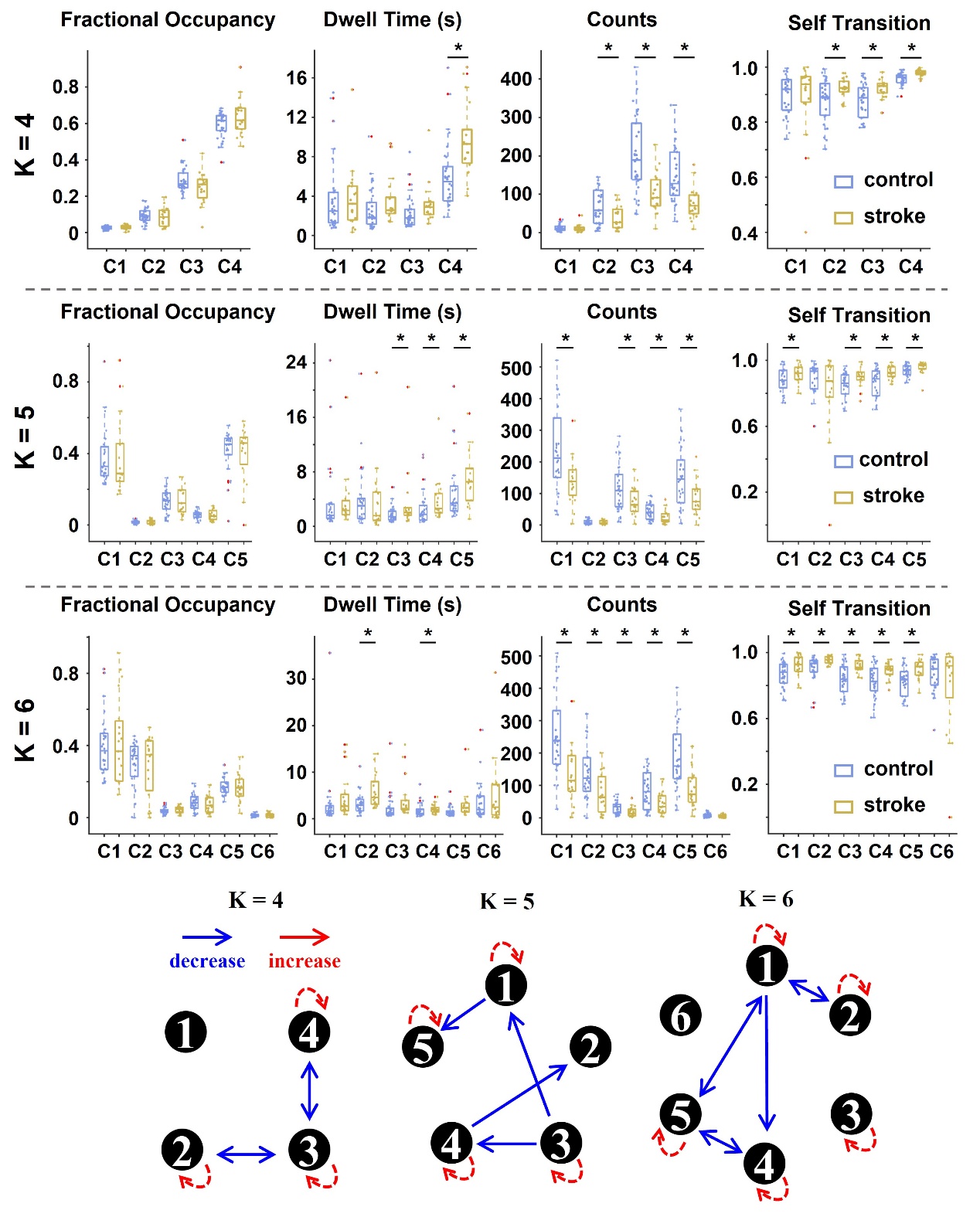

**Fig. S15** Stroke-induced impairment of functional state transitions in microcirculation. Across different cluster numbers, stroke increases the probability of remaining in individual functional states and prolongs their duration, while simultaneously reducing both transition probability and the total number of state transitions. Group differences between control (n = 30) and stroke (n = 19) are assessed using an unpaired t-test. *p < 0.05

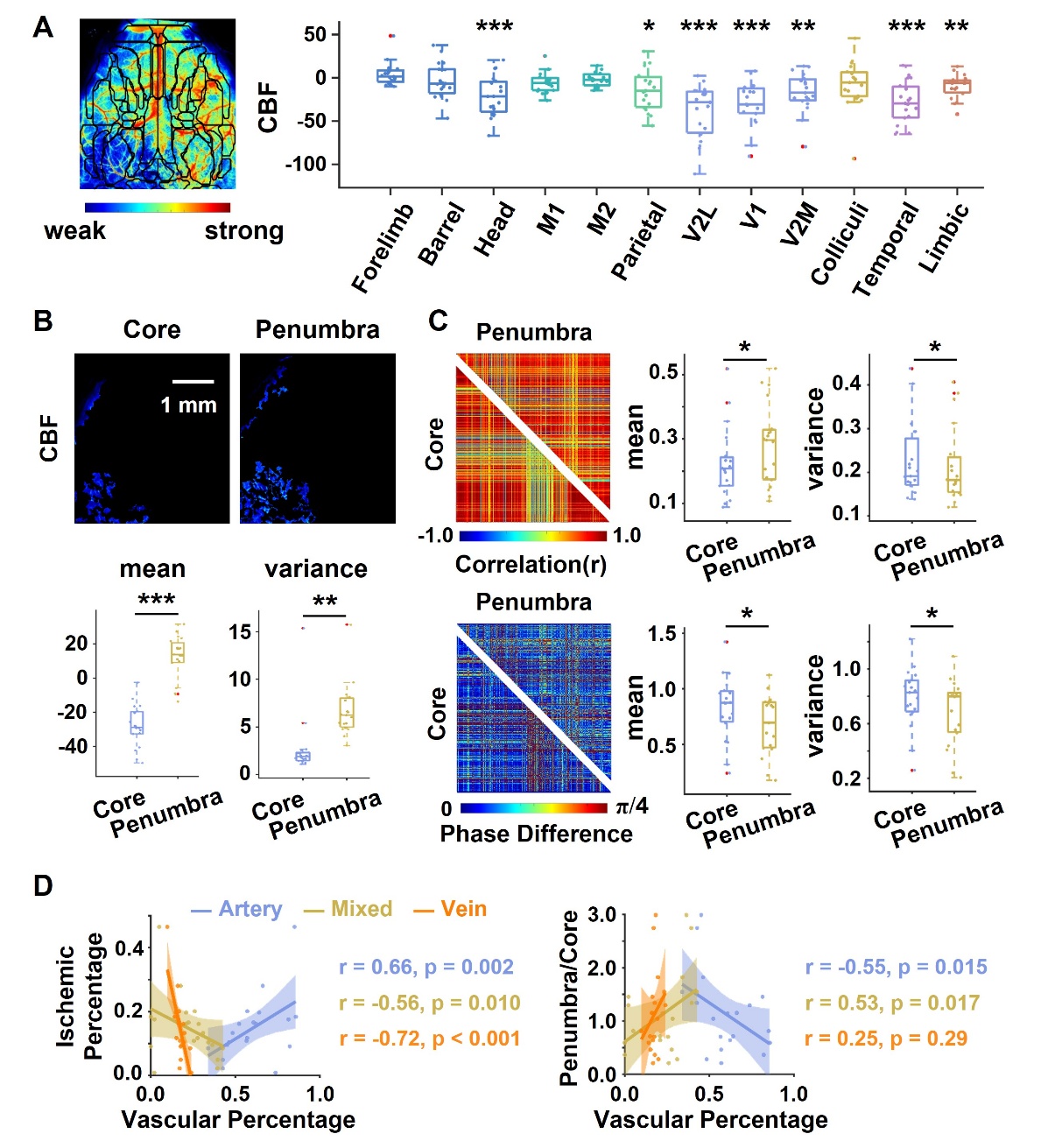

**Fig. 16** Quantitative profiling of cerebral blood flow impairment in ischemic regions. **A.** CBF intensity is lower in the left head, visual, temporal, and limbic regions than in the corresponding regions of the right hemisphere. Homologous differences between ischemic and contralateral hemispheres are analyzed with a one‑sample t‑test (n = 19). **B, C.** The ischemic penumbra exhibits preserved blood flow function relative to the impaired infarct core. CBF intensity and variability are significantly higher in the penumbra, accompanied by robust functional connectivity metrics, including higher mean inter-voxel correlation coefficients, smaller mean phase differences, and lower variance in functional connectivity matrices. Metric differences between the ischemic core and penumbra are assessed using a paired t-test (n = 19). **D.** Vascular architecture influences regional susceptibility to ischemic damage. Arterial vessel proportion correlates positively with ischemic lesion extent, whereas mixed and venous vessel proportion show negative correlations. Notably, the proportion of arteriovenous mixed region positively predicts the penumbra/core ratio. *p < 0.05, **p < 0.01, ***p < 0.001

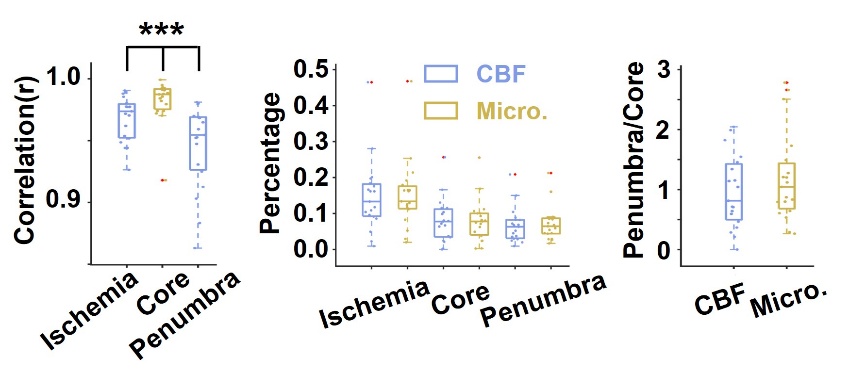

**Fig. 17** Comparative analysis revealed a strong correlation between ischemic territories delineated by CBF and those identified by microcirculatory assessment. Notably, the ischemic core showed greater agreement between the two methods than the penumbra regions. Despite this differential correlation pattern, the spatial extent of ischemic regions defined by either method did not differ significantly. Differences in ischemic territory metrics are evaluated via paired t-test with FDR adjustment (n = 19). ***p < 0.001
